# PI4P and BLOC-1 remodel endosomal membranes into tubules

**DOI:** 10.1101/2021.10.21.465321

**Authors:** Riddhi Atul Jani, Aurélie Di Cicco, Tal Keren-Kaplan, Silvia Vale-Costa, Daniel Hamaoui, Ilse Hurbain, Feng-Ching Tsai, Mathilde Dimarco, Anne-Sophie Macé, Yueyao Zhu, Maria João Amorim, Patricia Bassereau, Juan S. Bonifacino, Agathe Subtil, Michael S. Marks, Daniel Lévy, Graça Raposo, Cédric Delevoye

## Abstract

Intracellular trafficking is mediated by transport carriers that originate by membrane remodeling from donor organelles. Tubular carriers play major roles in the flux of membrane lipids and proteins to acceptor organelles. However, how lipids and proteins impose a tubular geometry on the carriers is incompletely understood. By exploiting imaging approaches at different scales on cells and in vitro membrane systems, we show that phosphatidylinositol-4-phosphate (PI4P) and biogenesis of lysosome-related organelles complex 1 (BLOC-1) govern the formation, stability and functions of recycling endosomal tubules. Endosomal PI4P production by type II PI4-kinases is needed to form nascent curved tubules through binding of BLOC-1 that stabilize and elongate them. Membrane remodeling by the PI4P/ BLOC-1 module functions not only in the recycling of endosomal cargoes, but also in the lifecycles of intracellular pathogens such as *Chlamydia* bacteria and influenza virus. This study demonstrates how a phospholipid and a protein complex coordinate as a minimal machinery to remodel cellular membranes into functional tubes.

## INTRODUCTION

The membrane-bound organelles of eukaryotic cells perform specific functions that are collectively needed for physiology. As a common feature, their limiting membrane isolates their lumen from the cytosol, while being actively remodeled to facilitate the exchange of components needed for the identity and the function of organelles (Jarsch et al., 2016). Such remodeling of the organellar membranes leads to the biogenesis of tubulo-vesicular carriers through the bending, invagination or evagination, and scission at precise membrane subdomains (McMahon and Gallop, 2005).

Relative to vesicles, tubular transport intermediates display a high surface-to-volume ratio that is best suited for transporting lipids and membrane-associated proteins (Maxfield and McGraw, 2004). Tubular intermediates consist of elongated curved membrane structures that are extend from largely planar membrane bilayers of many donor organelles [e.g. plasma membrane (PM), *trans*-Golgi Network (TGN), early endosomes, lysosomes and lysosome-related organelles (LROs)]. Their biogenesis and maintenance require membrane asymmetry imposed by extrinsic and/ or intrinsic factors (Stachowiak et al., 2013). Extrinsic factors include proteins that sense, impose, and/ or stabilize specific membrane curvatures [e.g., BAR (Bin-amphiphysin-Rvs)-domain containing proteins] and cytoskeletal elements that exert mechanical force onto membranes. Intrinsic factors include the accumulation of membrane components with a characteristic shape (e.g., conical lipids or proteins) that can form membrane domains with a particular geometry (Jarsch et al., 2016).

Among membrane lipid species, phosphatidylinositols (PtdIns or PI) are minor components of total cellular phospholipids but play crucial roles in membrane trafficking. Up to 7 different PI-Phosphate (PI_X_P) species can be generated, through reversible phosphorylation of their myo-inositol ring at positions -3, -4 and/ or -5. PI_X_Ps contribute to the identity and function of organelles by dynamically concentrating at membrane subdomains, where they aid in the recruitment and/or stabilization of specific proteins that modulate the membrane biophysical properties (Balla, 2013). For instance, PI(4,5)P_2_ at the PM cooperates with clathrin adaptor proteins to form carriers during endocytosis (Chang-Ileto et al., 2011). PI3P on early endosomes favors membrane binding of trafficking effectors (Gaullier et al., 2000; Xu et al., 2001) that support endosomal cargo recycling (Wallroth & Haucke, 2018), while PI4P (a.k.a., PtdIns4P) cooperates with membrane-shaping proteins and actin-based motors to effect the formation of TGN-derived carriers (Rahajeng et al., 2019). Thus, the local concentration of PI_X_P defines a membrane trafficking hotspot for membrane remodeling events during the biogenesis of transport carriers (Di Paolo and De Camilli, 2006; Saarikangas et al., 2010; Lemmon and Ferguson, 2000; Kutateladze, 2010).

Recycling endosomes (RE) are organelles that comprise a network of membranes tubules arising from the vacuolar domain of early sorting endosomes (SE) (Willingham et al., 1984; Yamashiro et al., 1984) and facilitating the sorting and trafficking of multiple cargoes to the PM and TGN, or to LROs in specialized cells (Delevoye et al., 2019; Klumperman and Raposo, 2014). Cargo transport through RE regulate many physiological processes at the cellular level such as nutrient uptake, cell migration, polarity, division or signaling, and autophagy, and at the tissue level such as neuronal plasticity and skin pigmentation (Grant and Donaldson, 2009; Puri et al., 2013; Delevoye et al., 2019). As such, RE malfunction is associated with diseases such as neurological disorders (e.g., Huntington and Alzheimer diseases) (Li et al., 2008; Zhang et al., 2006) or genetic forms of oculocutaneous albinism [e.g. Hermansky-Pudlak Syndrome (HPS)] (Bowman et al., 2019). In addition, RE are hijacked during certain viral and bacterial infections (Allgood and Neunuebel, 2018; Vale-Costa and Amorim, 2016). However, it remains to be determined which and how specific lipids and proteins coordinate to remodel endosomal membranes to generate and stabilize RE tubules.

While PI4P is enriched on Golgi membranes (Graham and Burd, 2011), a cohort of PI4P localizes also to early endosomes (Hammond et al., 2009) and endosome-derived vesicles (Ketel et al., 2016) or tubules (Jović et al., 2009). Cargo trafficking through the early endosomal system requires the spatiotemporal control of PI4P metabolism by two endosome-associated kinases (PI4KIIα and -β) (Craige et al., 2008; Wieffer et al., 2013; Hammond et al., 2014) and at least one phosphatase (Sac2) (Nakatsu et al., 2015; Hsu et al., 2015) to respectively synthesize and deplete PI4P on endosomal membranes. Loss of function or expression of either PI4KIIα or Sac2 results in missorting of the transferrin receptor (TfR) and the epidermal growth factor receptor (EGFR) (Minogue et al., 2006; Henmi et al., 2016; Nakatsu et al., 2015; Hsu et al., 2015), and PI4P contributes to stabilize some endosomal tubules (Jović et al., 2009). Moreover, loss of function of the PI4KII orthologue in *Drosophila melanogaster* larvae impairs RE function in the maturation of salivary gland secretory granules (Ma et al., 2020). However, how PI4P production and consumption regulate RE tubule biology is not understood.

The eight-subunit biogenesis of lysosome-related organelle complex 1 (BLOC-1) is required for the elongation and release of nascent RE tubular carriers from SE in coordination with the microtubule-based kinesin-3 motor KIF13A and actin-related machineries (Ripoll et al., 2016; Delevoye et al., 2014, 2016). Certain BLOC-1-dependent RE carriers deliver a subset of cargoes to LROs in specialized cell types, such as melanosomes in pigment cells, and thus loss of BLOC-1 function underlies several variants of HPS (Bowman et al., 2019). BLOC-1 physically associates with PI4KIIα (Salazar et al., 2009; Larimore et al., 2011), and thus may be integrated with PI4P functions at endosomal membranes. Here, we used a combination of biochemical approaches, light and electron microscopy (EM) on live and fixed cells, and *in vitro* cryo-EM imaging of model membranes to test whether BLOC-1 and PI4P orchestrate the biogenesis of RE tubules. We show *in vitro* that BLOC-1 binds to and tubulates PI4P-positive (^+^) membranes. In cells, endosomal PI4P contributes to remodel and stabilize membranes into RE tubules. The PI4P-asociated membrane remodeling functions allows for recycling of endosomal cargoes, and are exploited together whith BLOC-1 by the bacterium *Chlamydia* and by influenza A virus during their intracellular cycle. Our studies define a PI4P/ BLOC-1 module as a minimal machinery to generate and stabilize functional RE tubules, that is hijacked by certain infectious agents.

## RESULTS

### PI4P associates with recycling endosomal tubules

To detect RE tubules by fluorescence microscopy (FM), we performed live imaging of HeLa cells transiently transfected with plasmids encoding the kinesin-3 proteins KIF13A or KIF13B fused to fluorescent proteins [i.e., KIF13A-YFP or mCherry-KIF13B (Delevoye et al., 2014; Yamada et al., 2014; Delevoye and Goud, 2015)]. KIF13A is an effector of the RAB11 subfamily of small GTPases (Delevoye et al., 2014). KIF13A over-expression in HeLa cells generated mCherry-RAB11A-labeled RE tubules extending towards the cell periphery (**Fig. 1 A**, top panels, arrowheads). Its close homologue, KIF13B, was shown to localize to RAB5^+^ early endosomes in mouse embryonic fibroblasts (Kanai et al., 2014) or to RAB6A^+^ secretory vesicles in HeLa cells (Serra-Marques et al., 2020). We found that mCherry- KIF13B (pseudocolored in green) expression in HeLa cells also generated numerous and long GFP-RAB11A^+^ RE tubules (pseudocolored in red) (**Fig. 1 A**, bottom panels, arrowheads) that co-localized with KIF13A-YFP (**Fig. 1 B**, arrowheads), and that were closely apposed to RAB5^+^ early endosomes (**Fig. S1 A**, arrowhead). Thus, both KIF13A and KIF13B generate RE tubules.

**Figure 1.**
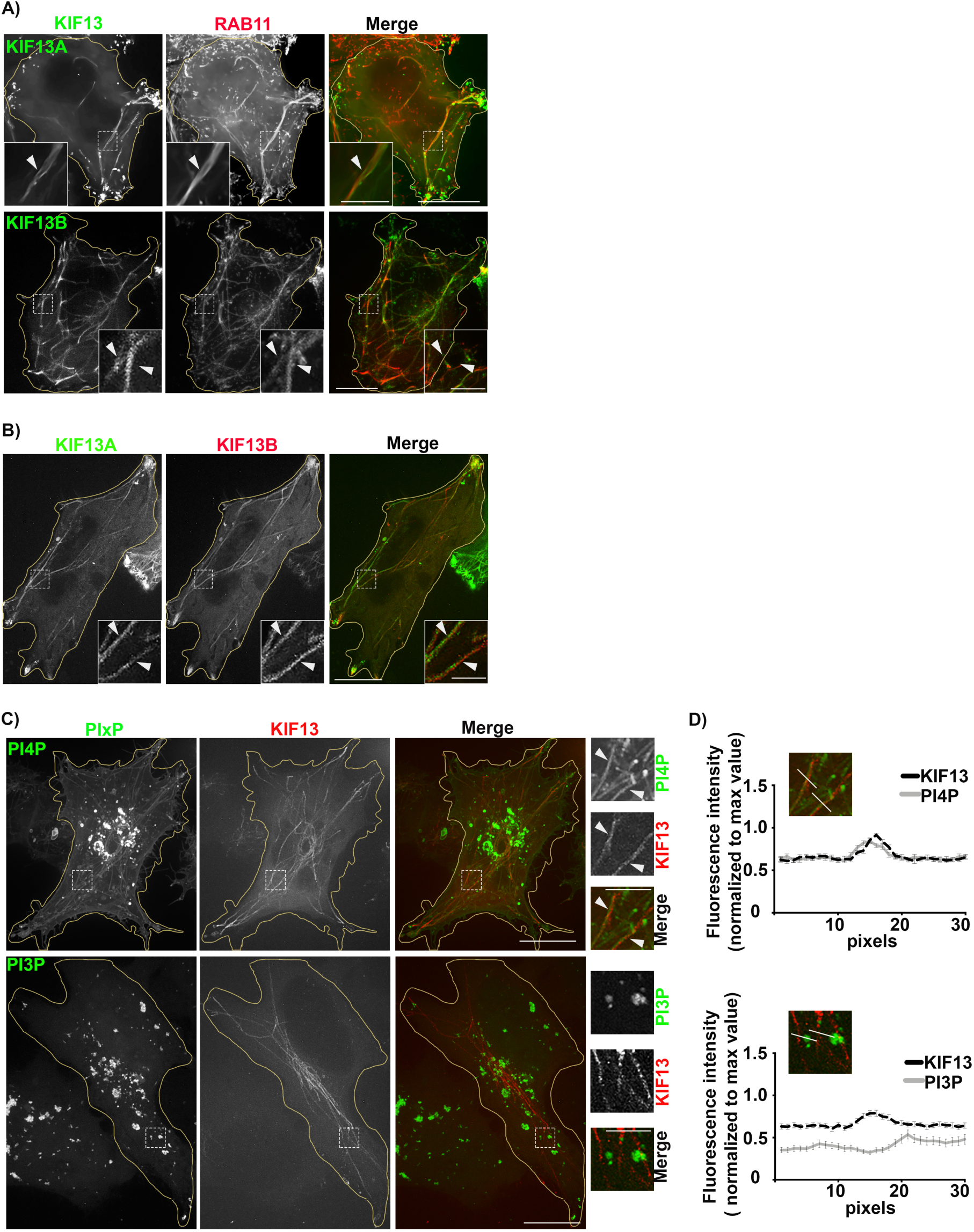
PI4P associates with KIF13^+^ recycling endosomal tubules. **A**) Live imaging frame of a Hela cell co-expressing KIF13A-YFP and mCh-RAB11A (green and red, respectively; top panels) or mCh-KIF13B and GFP-RAB11A (pseudocolored in green and in red, respectively; bottom panels). Magnified insets (4x) show RAB11A co- distribution with KIF13^+^ tubules (arrowheads). **B**) Live imaging frame of a Hela cell co- expressing KIF13A-YFP (green) and mCh-KIF13B (red). Magnified insets (4x) show KIF13A and KIF13B co-distribution (arrowheads). **C**) Live imaging frame of Hela cell expressing mCh-KIF13B (red) together with the GFP-coupled sensors (green) for either PI4P (SidC- GFP, top) or PI3P (GFP-2x-FYVE, bottom). Magnified insets (4x) show the co-distribution of KIF13B^+^ tubules with PI4P sensor (arrowheads), but not with the PI3P sensor. **D**) Line scan analyses intersecting KIF13B^+^ tubules captured as in C (=30 tubules per condition); values are mean ± SEM (n = 3 independent experiments). Cell periphery is delimited by yellow lines. Bars: (main panels) 10 µm; (magnified insets) 2.5 µm.

We next investigated the distribution of some PI_X_P species relative to RE tubules by co- expressing endosomal PI_X_P-specific fluorescent sensors with KIF13 proteins. Live-cell super-resolution FM (SR-FM; spinning-disk confocal microscopy using optical photon reassignment leading to 1.4 increased resolution) revealed that the PI4P sensor SidC-GFP largely localized to peripheral punctate structures devoid of iRFP-RAB5 (pseudocolored in red) (**Fig. S1 B**, top panels, arrowheads), and to large perinuclear structres that corresponded to the TGN labeled for TGN46 by immunofluorescence microscopy (IFM) (**Fig. S1 C**, top panels, arrowheads). The PI4P sensor also decorated to a lesser extent PM [here stained using fluorescently-conjugated wheat germ agglutinin (WGA); **Fig. S1 C**, bottom panels, arrowheads]. To analyze if a specific PI_X_P is enriched at RE tubules, we co- tranfected HeLa cells with plasmids encoding mCherry-KIF13B and sensors of PI4P (SidC-GFP) or PI3P (GFP-FYVE). Co-expression of mCherry-KIF13B led to the appearance of numerous PI4P^+^ tubular structures (**Fig. 1 C**, top panels), many of which co-distributed with mCherry-KIF13B (arrowheads) as shown by linescan analyses (**Fig. 1 D**, top panels). However, mCherry-KIF13B^+^ tubules were negative for the related PI3P sensor GFP-FYVE (**Fig. 1, C** and **D**, bottom panels), which instead largely labeled iRFP-RAB5^+^ early endosomes (**Fig. S1 B**, bottom panels, arrowheads). Together, these data indicated that KIF13 expression drives the generation of RE tubules with which PI4P is specifically and abundantly associated.

### BLOC-1 and PI4P generate tubules from membranes

Previously, we showed that the biogenesis of RE tubules from the limiting membrane of early endosomes relied on the BLOC-1-dependent elongation of short nascent tubes (Delevoye et al., 2016). Given that the PI4P sensor largely distributed to RE tubules and that BLOC-1 co-fractionated with the PI4P-producing kinase PI4KIIα (Salazar et al., 2009; Ryder et al., 2013), we hypothesized that PI4P and BLOC-1 cooperate to remodel the endosomal membrane during the formation of RE tubules. We first analyzed whether BLOC-1 interacted with membranes using a minimal system composed of purified BLOC-1 and lipid tubules and/ or vesicles. We expressed and purified the eight-subunits of BLOC-1 tagged with GST and 6xHis using a single polycistronic plasmid in *E. coli* [**Fig. S2 A**; and see (Lee et al., 2012)]. By negative staining EM, BLOC-1 appeared as curvilinear rods of ∼30 nm in length, and of variable curvatures (**Fig. 2 A**) as previously described (Lee et al., 2012).

**Figure 2.**
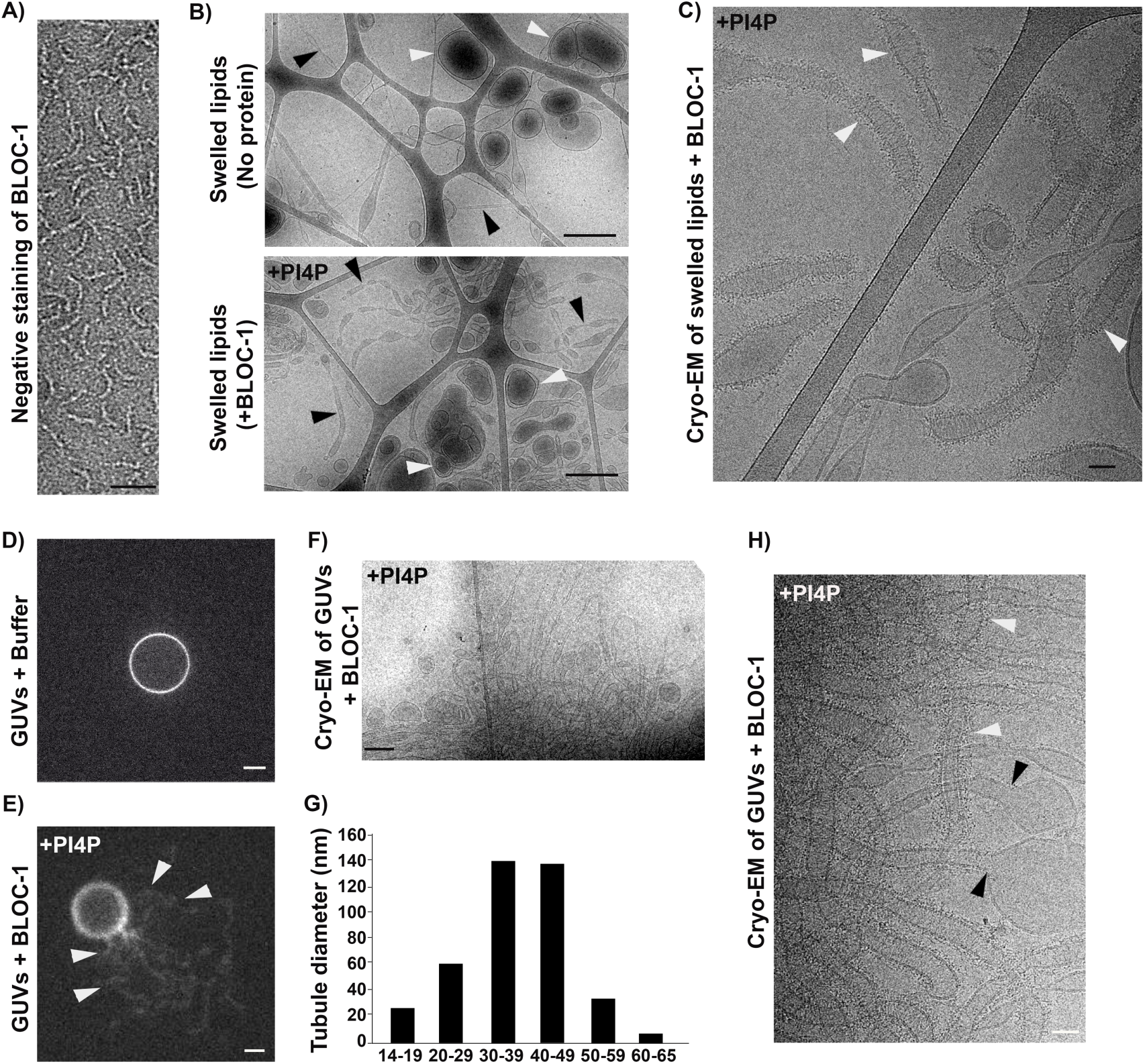
BLOC-1 in vitro generate tubules from PI4P containing membranes. **A**) Negatively stained image of BLOC-1 by EM. **B**) Cryo-EM images of a 50 μM resuspended lipid mixture of EPC/ bPS/ bPI4P (85/ 10/ 5 mol/mol) before (top ) or after (bottom) incubation with 0.16 μM of BLOC-1. Vesicles (white arrowheads) and tubules (black arrowheads) of different size and shapes were present. **C**) Cryo-EM images of tubule with BLOC-1 bound to membrane (white arrowhead). **D** and **E**) Fluorescent microscopy of EPC/ bPS/ bPI4P (85/ 10/ 5 mol/mol) GUVs before (**D**) or after (**E**) addition of BLOC-1, which led to the formation of tubules (arrowheads) at the surface of the GUV (see also **Supplementary Video 1**). **F**) Cryo-EM image of GUVs preparation shown in E revealing several protein-coated tubules. **G**) Plot of the diameter of the tubes generated with BLOC-1 from GUVs (n = 65). **H**) Cryo-EM images of tubules covered with BLOC-1 (white arrowheads) and of vesicles devoid of BLOC-1 (black arrowheads). Bars: (**A**) 25 nm, (**B**, and **F**) 500 nm, (**C** and **H**) 50 nm, (**D** and **E**) 5 μm. Figures are representative of at least three independent experiments.

We then analyzed the interaction of purified BLOC-1 with membranes containing different lipids. A membrane-bound lipid binding assay showed that BLOC-1-GST interacted with several mono- and di-phosphorylated PI_X_Ps, including PI4P (**Fig. S2 B**). We further analyzed the binding of BLOC-1 to lipid tubes and vesicles by cryo-EM. We prepared control lipid tubes of neutral charge made of galactocerebroside (GalCer)/ egg phosphatidylcholine (EPC) (**Fig. S2**, **C** and **D**; and see materials and methods). By cryo-EM imaging, GalCer/EPC tubes were of narrow and constant 27-nm diameter for lengths of several hundreds of nm (**Fig. S2**, **C** and **D**). Cryo-EM images of control lipid tubes incubated without (**Fig. S2 C**) or with BLOC-1 (**Fig. S2 D**) were similar and showed the two lipid leaflets of the tubes, but no BLOC-1 bound to the tubes (**Fig. S2 D**, arrowheads). By contrast, cryo-EM images of GalCer/ EPC tubes that were doped with either PI3P or PI4P (5 %) and incubated with BLOC-1 showed densities of black dots (arrrowheads) decorating the surface of both PI3P- (**Fig. S2 E**) and PI4P-doped (**Fig. S2**, **F** and **G**) tubes, showing that BLOC-1 binds to these negatively charged tubes.

We next investigated if BLOC-1 binds to PI4P^+^ membranes of larger curvature than the GalCer tubes by preparing lipid suspensions formed by swelling a dried lipid film containing EPC and 5 % PI4P. In the control condition without addition of BLOC-1 (**Fig 2 B**, top panel), tubules (black arrowheads) and vesicles (white arrowheads) of variable diameter were observed (8.5 tubules/ image, n = 20 images). Importantly, the addition of BLOC-1 (**Fig 2 B**, bottom panel) dramatically increased the number of tubules (black arrowheads; 18.6 tubules/ image, n = 23 images). At higher magnification by cryo-EM, BLOC-1 was present on tubules with diameters ranging from 25 nm to 80 nm (**Fig. 2 C** and **Fig. S2 H**, white arrowheads), but not on larger vesicles (**Fig. S2 H**, black arrowheads), suggesting that BLOC-1 tubulates membranes.

We next tested if BLOC-1 tubulates membranes by exploiting giant unilamellar vesicles (GUVs), which consist of a large reservoir of relatively flat and highly deformable membranes. Here, we used fluorescent GUVs containing 5 % PI4P with or without BLOC-1 (**Fig. 2**, **D-H**). While control GUVs (without addition of BLOC-1) appeared smooth and round by FM (**Fig. 2 D**), the addition of BLOC-1 induced the formation of tubules from the GUV surface (**Fig. 2 E**, arrowheads; and **Supplementary Video 1**). Cryo-EM analysis of the BLOC-1-treated GUVs (**Fig. 2 H** and **Fig. S2 I**) revealed that BLOC-1 decorated tubules with diameters of ∼40 nm (most varying between 20 and 60 nm) (**Fig. 2**, **G** and **H**; and **Fig. S2 I**, white arrowheads). The diameter of the tubules was not always constant (**Fig. 2 H** and **Fig. S2 I**). And, in most cases, tubules or vesicular buds derived from BLOC-1 swelled lipids or GUVs that are larger than 80 nm diameter were devoid of BLOC-1 (**Fig. 2 H** and **Fig. S2 H**, black arrowheads). Together, the results show *in vitro* that BLOC-1 transforms PI4P- containing vesicles into tubules with narrow but variable diameters by assembling and concentrating onto curved membranes.

### PI4KIIs are required for the formation and stabilization of recycling tubules

The *in vitro* data suggested that BLOC-1 could locally bind to PI4P^+^ and curved endosomal membranes prior to their elongation into tubules. We therefore investigated in cells if PI4P and its metabolism were needed for the biogenesis of RE tubules. Given that the production of PI4P on endosomal membranes relies on the PI4KIIα and PI4KIIβ (here after referred as PI4KII) (Hammond et al., 2014; Minogue et al., 2006; Balla and Balla, 2006), we examined the contribution of PI4KII to the formation of RE tubules by siRNA-mediated knock-down. HeLa cells treated with a combination of two individual siRNAs against PI4KIIα and PI4KIIβ (siPI4KII; see Materials and methods) showed ∼90 % reduction in both PI4KIIα and PI4KIIβ compared to control cells trated with non-targeting control siRNA (siCTRL) (**Fig. 3**, **A** and **B**; siPI4KIIα : 10.9 ± 0.4 %, siPI4KIIβ : 6.5 ± 3.5 %). By live-cell fluorescence imaging of siCTRL-treated cells transiently expressing KIF13A-YFP and mCherry-RAB11A, KIF13A and RAB11A largely decorated RE tubules extending, as expected, towards the cell periphery (**Fig. 3 C**, arrowheads; and **Fig. S3 A**, left panels). In contrast, tubules were largely absent in siPI4KII-treated cells, in which KIF13A-YFP and mCherry-RAB11A were instead localized to enlarged vesicular structures (**Fig. 3 C** and **Fig. S3 A**, right panels, arrows). Relative to control cells, the percentage of PI4KII-depleted cells that exhibited at least one KIF13A- YFP^+^ RE tubule was reduced by ∼30 % (**Fig. 3 D**; see materials and methods), and the RE tubules that were detected were ∼30 % shorter (**Fig. 3 E**; siCTRL: 2.2 ± 0.1 μm, siPI4KII: 1.5 ± 0.04 μm), revealing a defect in the generation and the elongation of RE tubules. This defect was specific to depletion of PI4KII, because siRNA-mediated depletion of the Golgi-associated PI4KIIIβ (Minogue, 2018) did not affect either the overall formation of KIF13A^+^ RE tubules or the percentage of cells displaying at least one RE tubule (**Fig. 3 F**, middle panel; and **Fig. 3**, **G** and **H**). The tubulation defect in PI4KII-depleted cells did not reflect an altered microtubule network, which was similar in siCTRL- and siPI4KII-treated cells (**Fig. S3 B**). Together, this set of experiments demonstrated that PI4KIIs are specifically required for the formation and/ or the elongation of KIF13^+^ recycling endosomal tubules.

**Figure 3.**
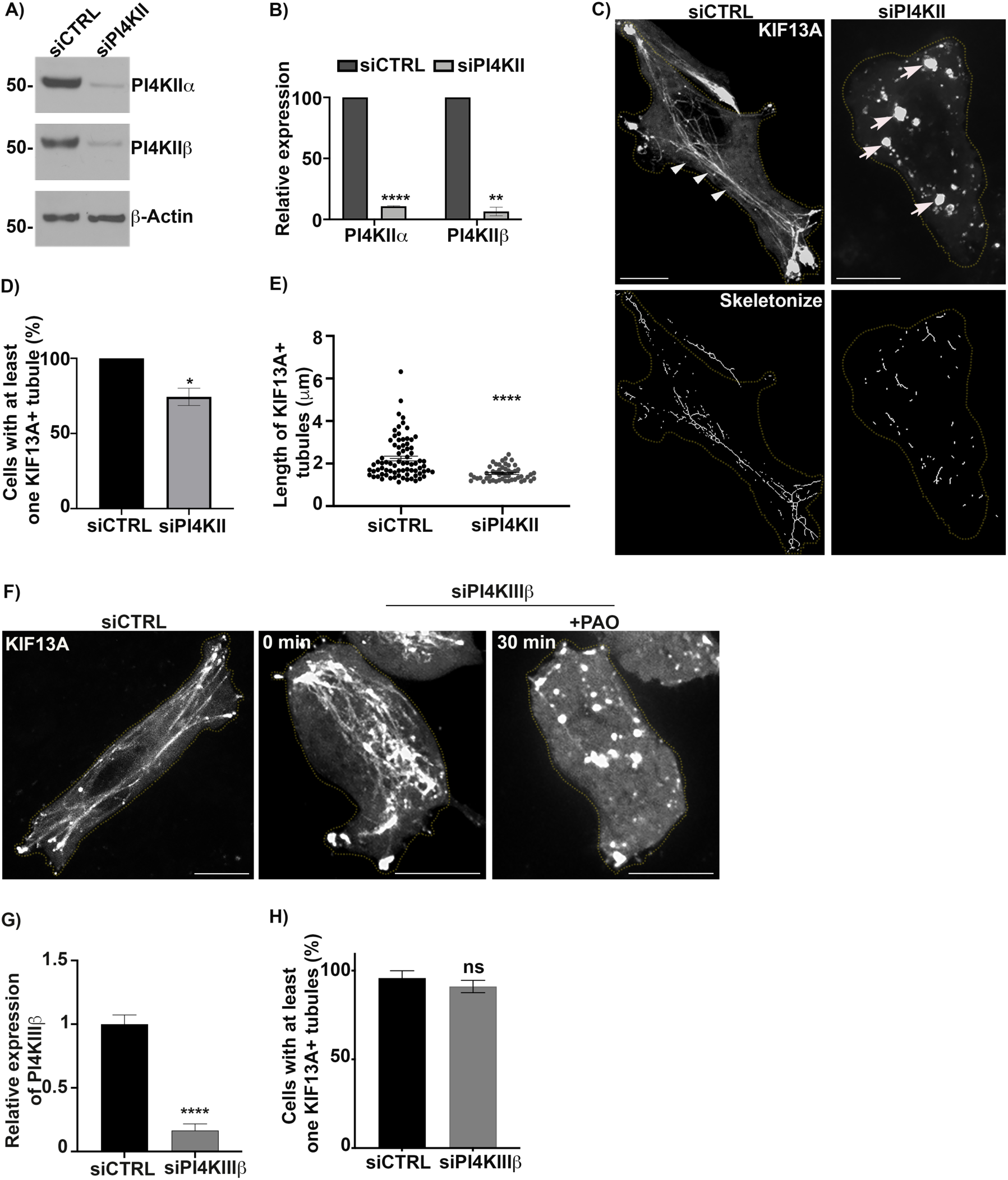
PI4KIIα and PI4KIIβ are required for the formation and stabilization of recycling endosomal tubules. **A**) Western blot of HeLa cell lysates treated with control (CTRL) or PI4KII (PI4KIIα and PI4KIIβ) siRNAs and probed for PI4KIIα (top), PI4KIIβ (middle) and β-Actin (loading control, bottom). **B**) Protein expression levels of PI4KIIα and PI4KIIβ in siCTRL and siPI4KII treated cells of 3 independent experiments were normalized to β-Actin levels. **C**) Live imaging frame (top) and associated binary images from “skeletonize” processing (bottom) of siCTRL- or siPI4KII-treated HeLa cells expressing KIF13A-YFP. Arrowheads, typical KIF13A^+^ RE tubules in siCTRL cells. Arrows, typical KIF13A^+^ vesicular structures in PI4KII-depleted cells. **D**) Quantification of the average percentage of siCTRL and siPI4KII cells (n > 60) in which at least one KIF13A^+^-YFP tubule was evident. **E**) Quantification of the average length of the KIF13A^+^-YFP tubules (n > 50 cells) in siCTRL and siPI4KII cells. **F**) Live imaging frame of KIF13A-YFP expressing siCTRL (left) or siPI4KIIIβ (middle and right) HeLa cells. Right panel shows siPI4KIIIβ-treated cells treated with PAO (600 nM, 30 min). **G**) Quantification of the PI4KIIIβ mRNA expression levels in siCTRL and siPI4KIIIβ cells by quantitative RT-PCR analysis relative to GAPDH as a reference gene. **H**) Quantification of the average percentage of siCTRL or siPI4KIIIβ-treated cells (n > 60) in which at least one KIF13A^+^-YFP tubule was evident. Cell periphery is delimited by yellow dashed lines. Data represent the average of at least three independent experiments and are presented as mean ± SEM. ns, not significant; *p < 0.05; **p < 0.01; ****p < 0.0001. Bars: 10 μm.

To assess if the enzymatic activity of the PI4KIIs was required to generate RE tubules, we tested the effect of phenylarsine oxide (PAO), a PI4KII inhibitor that causes cellular PI4P depletion (Yue et al., 2001). Similarly to PI4KII depletion, treatment of HeLa cells transiently expressing KIF13A-YFP for 30 min with PAO at low concentration (300 nM) reduced the percentage of cells harboring at least one KIF13^+^ tubule by ∼70%, as compared to cells treated with DMSO alone or with the PI3K inhibitor wortmannin (**Fig. S3**, **C** and **D**). Live imaging during the 30 min of PAO treatment showed that pre-existing KIF13A-YFP^+^ RE tubules were gradually lost, concomitant with the appearance of vesicular structures (**Fig. S3 E**, left panels), while the microtubule network remained intact (right panels). Finally, PI4KIIIβ- depleted cells similarly treated with PAO also displayed a loss of the KIF13A-YFP^+^ RE tubules (**Fig. 3 F**, right panel), showing that the stability of pre-existing RE tubules required specifically PI4KII enzymatic activity. Taken together, these data indicate that the formation of RE tubules and their stabilization rely on cellular PI4P synthesized by PI4KII.

### Endosomal PI4P stabilizes the recycling endosomal tubule

As PI4P is distributed among different organelles, including the Golgi apparatus and endosomes (Hammond et al., 2014), we asked which pool of PI4P contributes to generate the RE tubules. We exploited the rapamycin-inducible FRB-FKBP chemical dimerization system to trigger the recruitment of the FKBP-conjugated PI4P phosphatase enzymatic domain from Sac1 (FKBP-PJ-Sac) to early endosomal or Golgi membranes harboring FRB-conjugated RAB5 or -Giantin, respectively. Upon rapamycin addition, FKBP-PJ-Sac dimerizes rapidly with FRB-target proteins, allowing acute PI4P dephosphorylation on the cytosolic leaflet of the target organelle [**Fig. 4 A**; and (Hammond et al., 2014)]. HeLa cells expressing mRFP-FKBP-PJ-Sac, KIF13A-YFP, and either iRFP-FRB-RAB5 (**Fig. 4 B**) or FRB-Giantin (untagged; **Fig. 4 C**) were analyzed by live cell imaging before and after 20 min of rapamycin treatment. As expected, rapamycin addition led to the recruitment of PJ-Sac from a primarily cytosolic localization to RAB5^+^ early endosomes or the Giantin^+^ Golgi apparatus (**Fig. 4**, **B** and **C**, bottom left panels, arrows). FKBP-PJ-Sac recruitment to Golgi membranes had no impact on the appearance, number, or length of KIF13A-YFP+ RE tubules (**Fig. 4 C**, bottom right panel; and **Fig. 4**, **D** and **E**). However, FKBP-PJ-Sac recruitment to early endosomes had a dramatic effect (**Fig. 4 B**, bottom panels, arrows). The KIF13A-YFP^+^ RE tubules were reduced in both number (**Fig. 4 D**) and length (**Fig. 4E**), as had been observed in PI4KII-depleted or PAO-treated cells (**Fig. 3 C** and **Fig. S3 C**). Most of the KIF13A-YFP^+^ RE tubules in cells with early endosomal FKBP-PJ-Sac were suppressed and replaced by KIF13A-YFP^+^ vesicles, which overlapped with RAB5 and PJ-Sac signals (**Fig. 4 B**, arrows). These data support the conclusion that the stabilization of RE tubules requires the spatiotemporal control of PI4P levels on early endosomal membranes. Hence, together with the results presented in **Fig. 3**, we conclude that the pool of PI4P on early endosomal membranes functions in both the initiation and stabilization of RE tubules.

**Figure 4.**
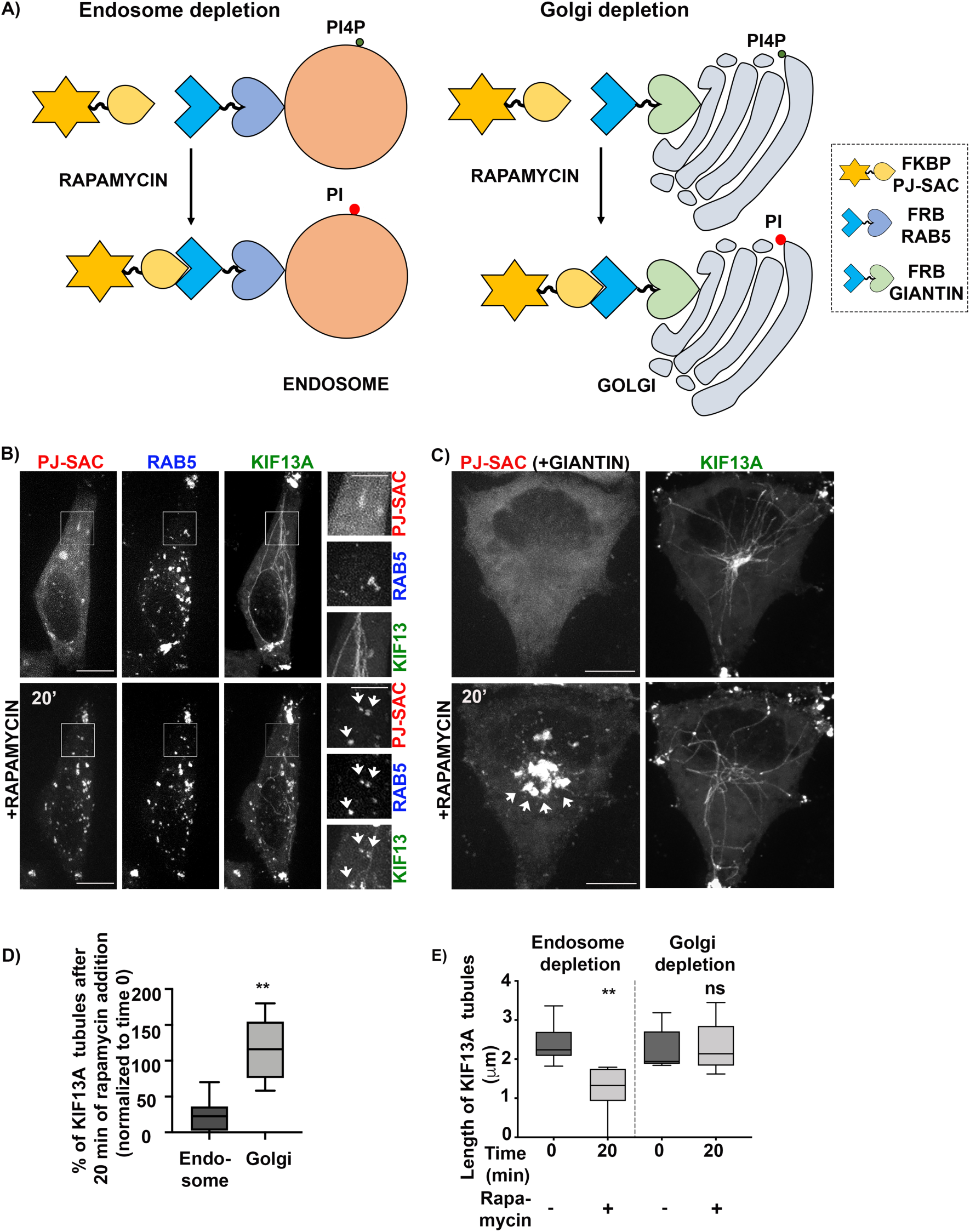
Deprivation of endosomal PI4P destabilizes recycling endosomal tubules. **A**) Schematic of the rapamycin-induced FRB-FKBP system allowing the organelle-specific depletion of PI4P through the recruitment of PI4P phosphatase Sac domain (mRFP-FKBP-PJ-SAC) to membranes positive for RAB5; (iRFP-FRB-RAB5, early sorting endosomes; left) or Giantin (FRB-Giantin, Golgi apparatus; right). The recruited Sac domain catalyzes the removal of phosphate (PO_4_) from PI4P (green ball) to generate PI (red ball). **B** and **C**) Live imaging frames of HeLa cells co-expressing KIF13A-YFP together with (B) mRFP-FKBP-PJ-Sac and either iRFP-FRB-RAB5 or (C) FRB-Giantin before (top) or after (bottom) addition of rapamycin (1 μM) for 20 min to recruit PJ-SAC to either RAB5^+^ endosomal (B) or Giantin^+^ Golgi (C) membranes (arrows). Note that acute targeting of PJ-SAC to RAB5^+^ membranes (B), but not to Giantin^+^ membranes (C), destabilizes the KIF13^+^ RE tubules. The FRB-Giantin chimera is not fluorescently-tagged, and thus is not imaged. **D**) Quantification of the average percentage of KIF13A-YFP^+^ RE tubules remaining 20 min after the addition of rapamycin relative to time 0 min in cells treated as in B (left) or in C (right). **E**) Quantification of the average length (μm) of KIF13^+^ RE tubule before (0 min) and 20 min after addition of rapamycin in cells treated as in B (left) or in C (right). Data are presented as box-plot and represent at least five independent experiments. ns, non-significant. ** p < 0.01. D-E: two- tailed unpaired *t*-test; endosome depletion, n = 85 tubules; Golgi depletion, n = 118 tubules; from at least 5 cells per condition. Bars: (main panels) 10 μm; (insets) 2.5 μm.

### PI4KIIs are required for the budding and elongation of early endosomal membranes

To form a RE tubule, the early endosomal membrane must be locally remodeled to generate a nascent tubule that is then elongated and ultimately released by membrane scission (Delevoye et al., 2014, 2016). We thus investigated if an imbalance of PI4P metabolism affects the remodeling of early endosomal membranes. We performed conventional EM in PI4KII-depleted cells immobilized by high-pressure freezing (HPF), which maintains the cellular membranes in a nearly native state (Hurbain et al., 2017). In siCTRL-treated cells, most early endosomes — defined as electron-lucent and round membrane-bound compartments containing a variable number of intra-luminal vesicles (ILVs) — harbored one or more nascent tubule profiles at their limiting membrane (**Fig. 5 A**, arrowheads; quantification in **Fig. 5 C**; nascent tubules/ endosome, siCTRL: 1.07 ± 0.17, siPI4KII: 0.36 ± 0.07), as reflected by their average length-to-width ratio (1.8 ± 0.2; quantified in **Fig. S4 C**). In PI4KII-depleted cells (**Fig. 5 B**), the limiting membranes of early endosomes harbored fewer deformations (arrowheads) as revealed by the ∼50 % reduction of nascent tubules per endosomal structure (**Fig. 5C**). This reduction did not reflect a defect in early endosome biogenesis, as early endosomes were frequently observed in siPI4KII-treated cells – although many contained fewer ILVs (**Fig. 5 B**, arrows), suggesting a potential impairment in endosomal maturation. This result shows that PI4KII expression is required for the initiation of nascent early endosomal tubules.

**Figure 5:**
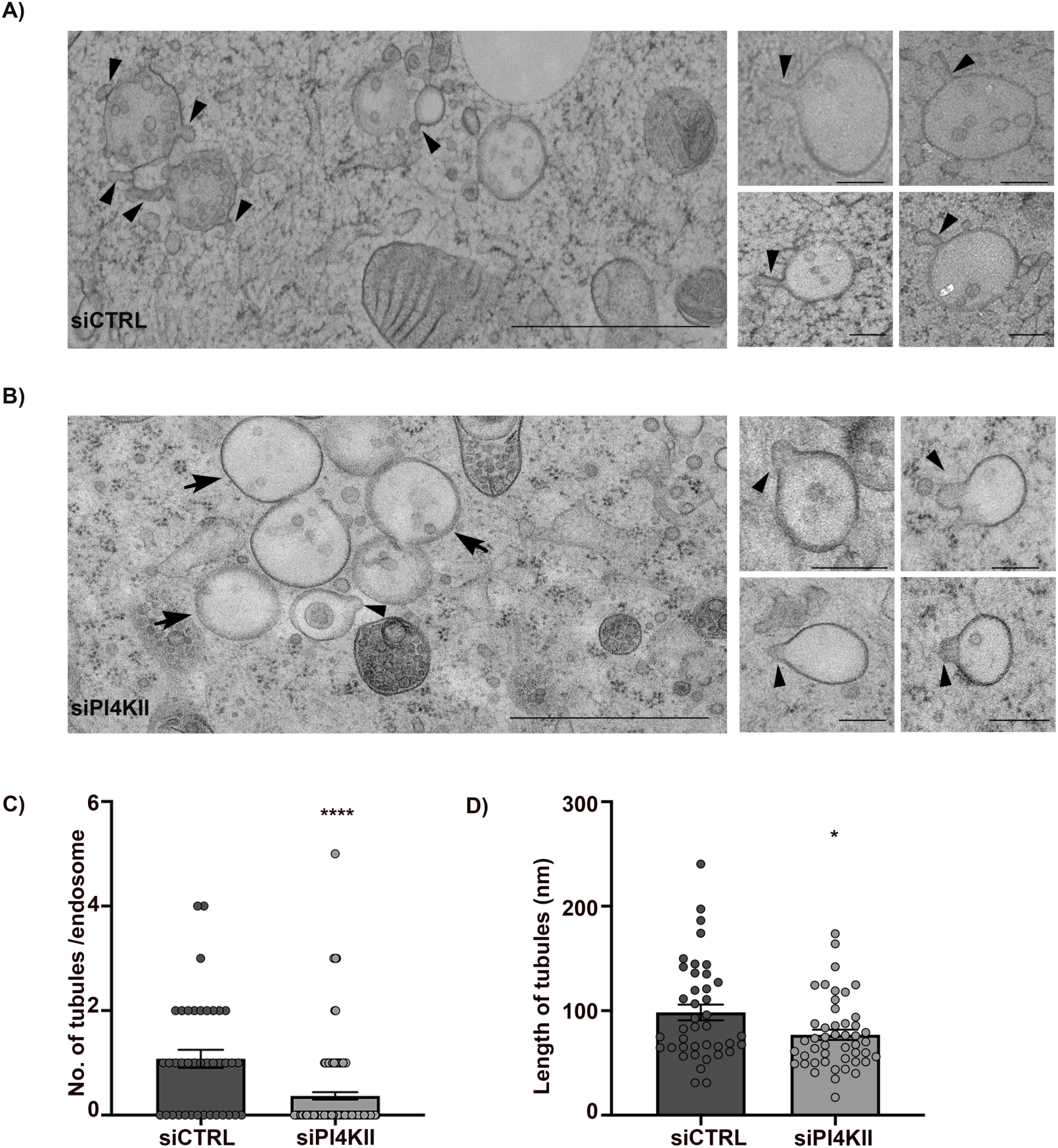
Depletion of PI4KIIs prevents the intiation of early endosomal tubules and their elongation. **A** and **B**) HeLa cells treated with control (siCTRL; A) or PI4KII siRNAs (siPIK4II; B) were immobilized by high-pressure freezing, and ultrathin sections were analyzed by conventional EM. Arrowheads point to nascent tubular structures emerging from the limiting membranes of endosomes. Arrows (B) point to endosomes without vesicular or tubular budded structure in PI4KII-depleted cells. Insets are additional representative endosomal structures captured in cells as in A and B. Arrowheads point to budded structures. **C**) Quantification of the average number of nascent tubules per endosome in siCTRL- and siPIK4II-treated HeLa cells as in A and B. **D**) Quantification of the average maximum length of the endosomal nascent tubules in siCTRL (n = 40) and siPI4KII (n = 47) treated HeLa cells identified in C. Data are the average of two independent experiments presented as the mean ± SEM (siCTRL, n = 8 cells; siPI4KII, n = 14 cells). C, D: two-tailed unpaired *t*-test. *p < 0.05; **p < 0.01; ****p < 0.0001. Bars, (main panel) 1 μm; (inset) 200 nm.

Though nascent tubules associated with endosomes were less numerous in PI4KII-depleted cells, three parameters were measured for those that were detected (**Fig. 5 B**, arrowheads; and **Fig. 5 C**): the maximum length, the maximum width, and the neck width (**Fig. S4 A**). In PI4KII-depleted cells, the nascent tubes were significantly shorter (**Fig. 5 D**; siCTRL: 98 ± 8 nm, siPI4KII: 76 ± 5 nm), consistent with a requirement for PI4KII and PI4P in tubule extension. The average maximum width of the nascent tubules was similar (**Fig. S4 B**, siCTRL: 60 ± 3 nm, siPI4KII: 61 ± 3 nm). Hence, the average length-to-width ratio of the nascent endosomal tubules in PI4KII-depleted cells was significantly reduced as compared to the control (**Fig. S4 C**; siCTRL: 1.8 ± 0.2, siPI4KII: 1.4 ± 0.1). Tubular membrane carriers constrict their neck prior to their release (Ripoll et al., 2018). The average neck width of the nascent tubules was marginally increased in PI4KII-depleted cells relative to control cells (**Fig. S4 D**; siCTRL: 49 ± 3 nm, siPI4KII: 58 ± 4 nm), although the increase did not reach statistical significance. These data indicate that PI4KII mainly controls the length of the nascent RE tubules.

Together, our data indicate that PI4KII and its PI4P product contribute to sequential steps – initiation and elongation – in the remodeling of the early endosome limiting membrane into a RE tubule .

### PI4KIIs control endosomal cargo recycling

If early endosomal PI4P contributes to RE tubule initiation, elongation and stabilization, then PI4KII depletion should impair the endosomal recycling of conventional cargoes. To test this prediction, we analyzed the dynamics of internalized transferrin (Tf) as a readout for the model recycling cargo, Tf receptor. Control- and PI4KII-depleted cells were pulsed for 10 min with fluorescent-Tf (Alexa Fluor 488 conjugated Tf) and then chased for different time points before processing by FM (**Fig. S5 A**) and quantification of signal intensity (**Fig. S5**, **B** and **C**). First, the overall Tf fluorescence intensity at time 0 of chase was ∼20 % reduced in PI4KII-depleted cells as compared to controls (**Fig. S5**, **A** and **B**), showing that less Tf was internalized during the pulse. Second, the overall intracellular distribution of Tf^+^ structures in PI4KII-depleted cells was distinct from the control cells (**Fig. S5 A**). At time 0, Tf^+^ endosomes in control- and PI4KII-depleted cells were distributed throughout the cell periphery (top panels). Whereas the general peripheral distribution tended to be maintained in control cells throughout the chase, Tf^+^ endosomes in PI4KII-depleted cells were more prominent in the perinuclear area after both 20 and 40 min of chase (middle and bottom panels, arrowheads). Third, whereas control cells showed a linear decay of the average normalized fluorescent intensity of Tf over the 40 min of chase, the decay of Tf intensity in PI4KII-depleted cells showed a significant plateau between the 20- and 40 min time points (**Fig. S5 C**), consistent with a recycling defect. All of these results were consistent with previous observations in cells expressing a catalytically inactive Sac2 mutant or Sac2 null cells (Hsu et al., 2015), and together show that PI4KII is required for the recycling of endosomal cargoes.

### PI4KII and BLOC-1 membrane functions are exploited by intracellular pathogens to enable their developmental cycle

Several intracellular pathogens exploit RE to establish their niche and/or to replicate. Thus, we reasoned that such pathogens might require PI4KII and BLOC-1 for certain steps of their developmental cycle. Among pathogen model systems, we chose 1) the influenza A virus (IAV), which exploits the RE-associated KIF13A and RAB11 to transport and propagate viral genome segments (viral ribonucleoproteins; vRNPs) during infection (Vale-Costa and Amorim, 2016; Ramos-Nascimento et al., 2017; Alenquer et al., 2019), and 2) the obligate intracellular bacteria *Chlamydia trachomatis* (*C. trachomatis*), which develops in a membrane-bound vacuole (referred to as an inclusion) from which tubules decorated by the RE-associated components RAB11, PI4KII and PI4P emerge (Rzomp et al., 2003; Moorhead et al., 2010), but whose function is unknown. The IAV model was used to define if PI4KII contributes to RE functions that are hijacked during viral infection, and the membrane tubules emanating from the *C. trachomatis* inclusion served as a proxy for PI4KII functions during membrane tubulation.

First, we explored the role of PI4KII in viral replication by IAV. In human A549 lung carcinoma cells, treatment with PI4KII siRNAs significantly reduced the expression of PI4KIIα and PI4KIIβ mRNAs (**Fig. S6**, **A** and **B**). Cells treated with siCTRL or siPI4KII were mock-infected or infected with IAV/Puerto Rico/8/34 (hereafter referred as PR8 virus; see materials and methods), and viral production was assayed. Importantly, PI4KII depletion led to a modest but consistent ∼10 % reduction of PR8 virus production at 8 hours post-infection (h.p.i.) relative to siCTRL-treated cells (**Fig. 6 A**). Given that the synthesis of viral matrix and capsid proteins was not affected by PI4KII depletion (**Fig. S6**, **C-E**), this finding suggested that the formation of IAV inclusions could be impaired. Viral inclusions are viewed as sites where complete genomes assemble before packaging into nascent virions to form fully infectious particles (Amorim, 2019). The packaging of influenza segmented genomes into productive virions requires RAB11, which redistributes during infection to intracellular sites of progeny RNA accumulation and is required to form liquid viral inclusions (i.e. not membrane bound), in which RAB11^+^ vesicles accumulate (Marta Alenquer et al., 2019). This is evident by IFM of PR8 infected cells, in which labeling for RAB11 and for IAV proteins such as NP overlap in punctate structures throughout the cell (Vale-Costa et al., 2016). Whereas both siCTRL- and siPI4KII-treated IAV-infected cells showed that RAB11^+^ puncta overlapped with NP labeling, these RAB11 and NP^+^ structures in PI4KII-depleted cells were substantially larger (**Fig. 6**, **B** and **C**). This difference in size could reflect aberrant viral inclusions with altered biophysical properties or a defect in transporting liquid viral inclusions. These data suggest that PI4KII contributes to the size and maintenance of liquid viral inclusions during IAV infection.

**Figure 6.**
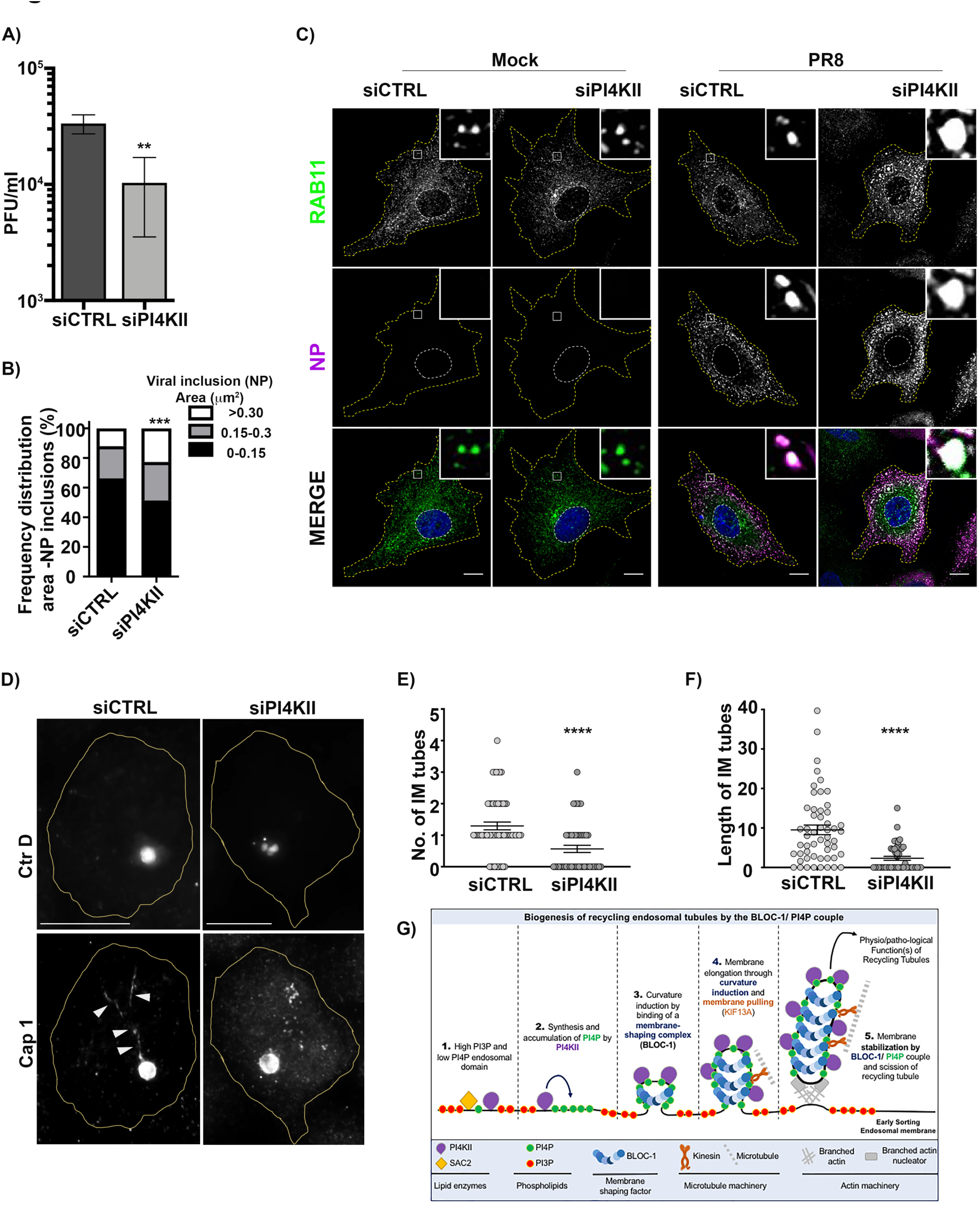
Intracellular pathogens exploit PI4KII and BLOC-1 to achieve their life cycle. **A**) Viral production assessed by plaque assay was plotted as plaque forming units per milliliter (PFU/ mL). A549 cells were treated with control siRNA (siCTRL) or siRNA specific for PI4KIIα and PI4KIIβ for 48h and then infected with PR8 at MOI 3 for 8h. A single experiment composed of 6 samples is shown and is representative of 3 independent experiments. Statistical analysis was performed using Student’s *t*-test (**p < 0.01). **B**) Frequency distribution of NP viral inclusions within the three area categories (in µm^2^) was plotted for each condition. **C**) Cells treated with control siRNA (siCTRL) or siRNA specific for PI4KIIα and PI4KIIβ and infected or mock-infected with PR8 at MOI 3 for 8h were processed by IFM. Cytosolic viral inclusions were identified using antibodies against RAB11 (green) and viral NP (magenta) proteins. Areas of RAB11 and NP co-localization are highlighted by white boxes. Nuclei (blue) and cell periphery are delimited by white and yellow dashed lines, respectively. **D**) HeLa cells treated with control (siCTRL) or PI4KII (siPI4KII) siRNAs were infected 12h with *Chlamydia trachomatis* serovar D *(Ctr D)* expressing mCherry (top panels), then fixed and analyzed by IFM using antibodies against the bacterial inclusion protein Cap1 (bottom panels). Arrowheads point to the Cap1^+^ tubules elongating from the inclusion membrane. Cell periphery is delimited by yellow lines. **E** and **F**) *Ctr D***-**infected cells were analyzed for the average number (E) and for the mean length (F) of Cap1^+^ tubules emanating from the bacterial inclusion as in D. **G**) Working model of the role of PI4P and PI4KII and BLOC-1 during the formation of recycling endosomal tubules from the early sorting endosomal membrane. (**1**) At the PI3P^+^ early endosomal membrane subdomain, PI4P locally produced by PI4KII is quickly depleted by SAC2 phosphatase activity. (**2**) In PI3P-deficient domains, endosomal PI4P locally synthesized by PI4KII could accumulate. (**3**) BLOC-1 binding to PI4P^+^ membrane subdomains facilitate the formation of buds or nascent tubules with diameter and curvature compatible with BLOC-1 association/ stabilization to membrane. (**4**) Nascent tubules accumulate PI4P produced by associated PI4KII that sustain elongation through continuous recruitment of BLOC-1 to PI4P+ tubular membrane and concomitant pulling by KIF13 motors along microtubules (Delevoye et al., 2016). (5) Eventually, nascent extended RE tubules are severed at their neck though BLOC-1 cooperation with actin-polymerizing machinery (Delevoye et al., 2016) to release functional PI4P^+^ tubules required for endosomal cargo (e.g. Tf) recycling. The machineries required for the biogenesis of recycling endosomal tubules can be exploited by pathogens to either use recycling endosomal membranes (e.g., IAV) or tubulate parasitaphorous vacuoles (e.g., Chlamydia). Statistical analysis of data (B) was performed using a non-parametric two-way ANOVA test, followed by Sidak’s multiple comparisons test (***p < 0.001). Approximately 45 cells were analyzed per condition and 3 independent experiments were performed. Data (E and F) are the average of three independent experiments presented as the mean ± SEM. E and F, two-tailed unpaired *t*-test (****p < 0.0001). Bars: 10 μm.

Second, we investigated the impact of PI4KII on the formation of membrane tubules from the surface of bacterial inclusions formed during *Chlamydia trachomatis* serovar D (Ctr D) infection of HeLa cells. CTRL- or PI4KII-depleted cells were infected with Ctr D expressing mCherry for 20 hours before analysis by IFM. In both controls and cells depleted of PI4KIIα and PI4KIIβ, Ctr D (**Fig. 6 D**, top panels) were internalized and developed within an inclusion (here detected using an antibody to the Cap1 inclusion membrane protein) of comparable size and shape (**Fig. 6 D**, bottom panels). However, relative to control cells, PI4KII-depleted cells harbored ∼60 % fewer elongated inclusion membrane (IM) tubes decorated by Cap1 (**Fig. 6 D**, arrowheads; and **Fig. 6 E**; IM tube/ cell, siCTRL: 1.3 ± 0.1; siPI4KII: 0.6 ± 0.1). Moreover, the remaining IM tubes in PI4KII-depleted cells were ∼70 % shorter than in controls (**Fig. 6 F**; siCTRL: 9.5 ± 1.2 μm, siPI4KII: 2.3 ± 0.5 μm). Similar results were observed in Ctr D infected cells that had been depleted of KIF13A or of BLOC-1 (**Fig. S6**, **F**- **H**). Compared to control cells, in PI4KII-depleted cells the IM tubes per cell were fewer in number (**Fig. S6 G;** siKIF13A: 0.8 ± 0.1, siBLOC-1: 0.8 ± 0.1) and shorter in length (**Fig. S6 H**; siKIF13A: 5.1 ± 1.0 μm, siBLOC-1: 5.8 ± 1.3 μm). These data revealed that components required to form RE tubules are also used to form tubules from the *Chlamydia* inclusion membrane.

## DISCUSSION

Recycling endosomal tubules are critical in the trafficking of essential physiological cargoes from early endosomes to other intracellular compartments. Here, by exploring the way lipids and proteins shape the early endosomal membrane to form RE tubules, we identified the phospholipid PI4P and the protein complex BLOC-1 as a unit that tubulates endosomal membranes. *In vitro* analysis with a minimal set of lipids shows that BLOC-1 binding to PI4P^+^ membranes generates tubules. As a working model (**Fig. 6 G**), we propose that the PI phosphatase Sac2 activity would maintain a low level of PI4P at the early sorting endosomal membrane (rich in PI3P) by counteracting the activity of PI4KIIs. The local elevation of PI4P concentration by PI4KIIs would favor BLOC-1 association to the endosomal membrane, which would further elongate through the combination of curvature induction and microtubule-based motor engagement [e.g., KIF13A (Delevoye et al., 2014)]. While the RE tubule is stabilized at least in part by the presence of BLOC1 bound to PI4P, BLOC-1 together with branched actin filaments would contribute to the constriction of the RE tubule neck prior to its release by membrane fission (Delevoye et al., 2016). Together, we conclude that BLOC-1 and PI4P play a central role in tubulogenesis during the initiation, elongation and stabilization of RE tubules.

PI4P metabolism relies on specific kinases (PI4KII and -III, each comprising two members, α and β) and phosphatases (Sac1 and Sac2), which control PI4P abundance on the cytosolic leaflet of membranes of various organelles (e.g., Golgi apparatus, endosomes, and lysosomes) and transport carriers derived therefrom. Sac2 localization is restricted to early sorting vacuolar endosomal domains (Nakatsu et al., 2015; Hsu et al., 2015), whereas PI4KIIα can also associate with derived endosomal tubules in Drosophila and mammalian cells (Burgess et al., 2012; Henmi et al., 2016; Minogue et al., 2006). This indicates that PI4P production and consumption at early endosomal membranes are differently balanced between the vacuolar and tubular domains. Indeed, we show that PI4P extensively decorates RE tubules bearing KIF13 and RAB11 (Delevoye et al., 2014). The relative PI4P abundance on RE tubules over RAB5^+^ early endosomes might thus reflect a prominent Sac2 activity at the sorting endosomal membrane that counteracts the PI4KII activity. Hence, a PI3P-to-PI4P gradient would be naturally implemented along the sorting-to-recycling pathway (Ketel et al., 2016). We show that the PI4P^+^/ KIF13^+^ RE tubules are apposed to PI3P^+^/ RAB5^+^ sorting endosomes, and that the hydrolysis of endosomal PI4P (via endosomal targeting of Sac2) destabilizes the RE tubules. Therefore, these results suggest that PI3P^+^ sorting and PI4P^+^ recycling endosomal membranes may form a continuum, as previously observed in pigment cells (Dennis et al., 2015; Delevoye et al., 2009).

Along such a continuum, local production of PI4P at the sorting endosomal membrane might be a signal to initiate the formation of RE tubules. Depleting PI4KII expression abrogates the biogenesis of RE tubules and leads to the accumulation of early vacuolar endosomes with fewer nascent tubules. This suggests that PI4P must first be produced at the sorting endosomal membrane to recruit locally a curvature-inducing component, like BLOC-1, that contributes to initiate and/ or to stabilize membrane curvature prior to elongation to form a RE tubular carrier. The PI4P contribution to the biogenesis of tubular transport carriers might not be restricted to endosomal membranes, as PI4P together with curvature-inducing proteins could aid their formation from various membrane sources, including the Golgi apparatus (Dippold et al., 2009; Rahajeng et al., 2019), maturing secretory granules (Ma et al., 2020) autolysosomes (McGrath et al., 2021), or phagolysosomes (Levin-Konigsberg et al., 2019; López-Haber et al., 2020).

We show that acute inhibition of PI4KII activity or rapid hydrolysis of endosomal PI4P destabilize pre-existing RE tubules. Moreover, the few tubules that still emerged from vacuolar sorting endosomes in cells depleted of PI4KII were unusually short, suggesting that PI4P contributes not only to the initiation of RE tubules, but also to their elongation and/or stabilization. The elongating/stabilizing function might not be limited to KIF13^+^ RE tubules, as PI4P decorates and maintains other extended endosomal tubules with recycling functions (Jović et al., 2009). Given that PI4P, together with BLOC-1, contributes first to initiate the RE tubule and second to stabilize it, this implies that one or more PI4Ks must be required to supply PI4P along the tubules. This is consistent with data in melanocytes that both PI4KIIα and PI4KIIβ are present on melanosome-bound RE tubules and that both are required for their stability (Zhu et al., companion paper). Because we show that the depletion of PI4KIIIβ, which interacts with RAB11 (Burke et al., 2014), does not affect the formation or stability of KIF13^+^ RE tubules — except when PI4KII enzymatic activity is concomitantly inhibited — we propose that only PI4KII (α and/ or β) produce PI4P (*i*) at the early sorting endosomal membrane to facilitate the biogenesis of RE tubules, and (*ii*) along the RE tubules to stabilize them.

We show that BLOC-1 can bind *in vitro* to model membranes containing PI4P. The eight-subunit BLOC-1 cooperates with KIF13A to elongate nascent RE tubules (Delevoye et al., 2016), with which it associates (Di Pietro et al., 2006). By negative stain EM imaging, purified recombinant BLOC-1 consists of a 30 nm long and flexible curvilinear bow-shaped protein complex (Lee et al., 2012), suggesting that BLOC-1 might adapt to membranes of variable diameters. Here, BLOC-1 tubulates relatively curved (i.e., small liposomes) or flat (i.e., GUVs) membranes provided they contain PI4P. In cells, it can be anticipated that locally raising the concentration of BLOC-1 on PI4P^+^ membranes would be sufficient to form and/or stabilize tubules. This is consistent with observations in cells, in which depleting BLOC-1 expression led to the accumulation of early sorting endosomes that harbored numerous buds that did not elongate (Delevoye et al., 2016). We therefore propose that BLOC-1 is a membrane shaping complex (Johannes et al., 2014) that facilitates tubulation of PI4P^+^ membranes.

Several membrane remodeling components form supramolecular assemblies on membranes, such as dynamin (Kong et al., 2018), the N-BAR protein Endophilin (Mim et al., 2012), ESCRT-III complex proteins (Nguyen et al., 2020), retromer (Kovtun et al., 2018) and COP-II complexes (Hutchings et al., 2021). Interestingly, our data suggest that BLOC-1 can be densely packed on PI4P^+^ nanotubes without forming a periodic arrangement. Given its 30 nm-length (Lee et al., 2012), several BLOC-1 molecules are likely needed to surround tubules. By organizing into various architectures, BLOC-1 assemblies could then stabilize tubules of different curvatures (and hence different diameters from 20-60 nm). Indeed, as shown by EM of purified BLOC-1, the linear chain made by the eight subunits can bend as much as 45° (Lee et al., 2012), suggesting together that BLOC-1 may tune its 3D architecture and/or assembly of molecules on curved membrane.

Since PI4P and BLOC-1 function as a module in endosomal membrane remodeling, loss of their functions should have similar consequences. As expected from previous studies (Henmi et al., 2016; Jović et al., 2009; Hsu et al., 2015) and as also observed in cells depleted of BLOC-1 or KIF13A (Delevoye et al., 2014; Setty et al., 2007), we show that cells depleted of PI4KII display slower Tf recycling associated with reduced Tf uptake, likely reflecting decreased Tf Receptor cell surface expression. BLOC-1-deficient cells also missort additional cargoes, particularly in melanocytes (Monis et al., 2017; Setty et al., 2007; Di Pietro et al., 2006; Delevoye et al., 2016), and consistently Zhu et al document missorting of similar cargoes in cells depleted of PI4KIIα or β (Zhu et al., companion paper). Thus, by initiating RE tubulogenesis, BLOC-1 and PI4KII together maintain the trafficking and sorting of cargoes along the endocytic pathway.

The requirement for PI4KII in RE biogenesis was validated in our analyses of PI4KII function in the developmental cycle of two intracellular pathogens. We show that the bacterium *Chlamydia trachomatis* uses PI4KII (as well as BLOC-1 and KIF13A) to build the parasitophorous inclusion-derived tubules, while the virus influenza A used PI4KII to form adequately sized IAV liquid viral inclusions and to control viral progeny production. The role of PI4KII in *Chlamydia* inclusion maturation is consistent with reports showing the recruitment of RAB11 (Rzomp et al., 2003) and of PI4KIIα, PI4P or PI4P-binding proteins (Moorhead et al., 2010) to the inclusion. Such redistribution of components allows the internalized bacteria to manipulate intracellular pathways, like the slow recycling endosomal pathway (Ouellette and Carabeo, 2010), and to complete the *Chlamydia* infectious cycle (Mölleken and Hegemann, 2017; Leiva et al., 2013). The role of PI4KII in IAV assembly is also consistent with reports of the exploitation of RE function by IAV. Cells infected by IAV redistribute RAB11 to vRNPs^+^ liquid structures, which relocalize to the cell periphery upon KIF13A overexpression (Vale-Costa and Amorim, 2016; Ramos-Nascimento et al., 2017; Amorim et al., 2011; Vale-Costa and Amorim, 2017; Alenquer et al., 2019). Given the requirement of PI4P and BLOC-1 during the building of functional RE tubules, it is likely that many other intracellular pathogens exploit these components to hijack the host RE system to their own benefit (Yong et al., 2021).

The dysregulation of PI4KII expression or genetic mutations of BLOC-1 subunits are associated with risks of development of cancers or neurological disorders (Minogue, 2018; Hartwig et al., 2018), and inactivating BLOC-1 subunit mutations are found in subtypes of HPS (Bowman et al., 2019), an inherited syndromic disorder characterized by albinism, excessive bleeding, and other defects. Given that an altered biogenesis and/ or function of RE has been characterized in these different disorders (Delevoye et al., 2019; O’Sullivan and Lindsay, 2020), a more systematic investigation of whether and when PI4P and BLOC-1 act together as a universal and minimal machinery reshaping membranes into tubules will provide a better understanding of the mechanisms of membrane dynamics in physiology and of their malfunction in pathophysiology.

## Author contributions

R.A.J. and C.D. conceived the study; M.J.A., J.S.B., A.S., M.S.M., D.L., G.R. and C.D. contributed to fund the project; R.A.J., A.D.C, T.K.K., S.V-C., D.H., F-C. T., A.S., D.L., G.R. and C.D. designed the research; R.A.J., A.D.C., T.K.K., S.V-C., D.H., I.H., F-C. T., M.D. and D. L. performed the experiments; R.A.J., A.D.C., T.K.K., S.V-C., D.H., F-C. T., M.D. and D.L. collected the data; R.A.J., A.D.C., T.K.K., S.V-C., D.H., A-S. M., A.S., D.L. and C.D. analyzed the data; R.A.J., T.K.K., S.V-C., M.J.A., D.H., A.S., M.S.M., D.L., G.R. and C.D. interpreted the results; R.A.J. and C.D. wrote the manuscript with contribution from A.D.C., T.K.K., S.V-C., D.H., F-C.T., M.J.A., P.B., J.S.B., A.S., M.S.M., D.L., and G.R.. All authors contributed intellectual capital into the study and edited versions of the manuscript.

## Competing Interest

The authors declare no competing interest.

## Acknowledgments

This work was supported by National Institutes of Health grants R01 EY015625 (to M.S.M. and G.R.), GENESPOIR (to C.D.), Institut National de la Santé et de la Recherche Médicale (INSERM), Institut Curie, and the Centre National de la Recherche Scientifique (CNRS), and by the Intramural Program of NICHD, NIH (ZIA-HD001607, to J.S.B.). We thank the Cell and Tissue Imaging core facility (PICT IBiSA), Institut Curie, member of the French National Research Infrastructure France-BioImaging (ANR10-INBS-04). We thank CryoCapCell (France) for providing equipment (HPM Live µ) and expertise in high pressure freezing, as well as Alexis Canette from the IBPS (Institut de Biologie Paris-Seine) electron microscopy core facility with the support of Sorbonne-Université and CNRS. We also thank Drs. Bruno Antonny and Bruno Mesmin (Université Nice Sophia Antipolis, France) for insightful discussions. This work was supported by a grant from the LabEx Cell(n)Scale (ANR-11- LABX-0038, ANR-10-IDEX-0001-02). M.J.A. is funded by The European Research Council consolidator grant under the European Union’s Horizon 2020 research and innovation program (No 101001521 - LOFlu) and by the Portuguese Fundação para a Ciência e a Tecnologia (CEECIND/02373/2020). S.V-C. is funded by a Junior Researcher working contract from Fundação para a Ciência e a Tecnologia (FCT) and Instituto Gulbenkian de Ciência (IGC, Portugal).

## Materials and Methods

### Cell culture, transfections, and infections

#### Cell Culture

Hela cells were grown in DMEM with GlutaMAX^TM^ (Invitrogen), supplemented with 10% FBS at 37°C under 5% CO_2_. For viral infection, the epithelial cells Madin-Darby Canine Kidney (MDCK) and A549 human alveolar basal carcinoma were kind gift of Prof Paul Digard (Roslin Institute, UK), cultured as previously described (Amorim et al., 2011), and regularly tested for mycoplasma contamination with the LookOut mycoplasma PCR detection kit (Sigma, MP0035), using JumpStart Taq DNA Polymerase (Sigma, D9307).

#### Transfection

For plasmid transfections in 6-well plate, cells [2.5 × 10^5^/ well or fluorodish (World Precision Instrument)] were seeded at day 1, transfected on day 2 with respective plasmids (300 ng – 1000 ng) using jetPRIME reagent (Polyplus-transfection), and processed on day 3 for IFM, live cell imaging or immunoblotting (IB). For rapamycin-inducible-sac system, live cell imaging was carried out by co-trasnfecting plasmids coding for mRFP-PJ-Sac-FKBP (600 ng) and KIF13A-YFP (1μg) together with FRB-ECFP(W66A)-Giantin (400 ng) or iRFP-FRB-RAB5 (400 ng) and imaging was carried out before and after addition of rapamycin (1 mM, 20 min) [see (Hammond et al., 2014) for details]. For siRNA transfections (HeLa cells), cells were seeded in 6-well plate on day 1 (1.5 × 10^5^/ well), transfected with respective siRNAs (200 pmol/ well) on day 2 and day 4 using oligofectamine (Invitrogen; 10 μl/ well), and processed on day 6 for IFM or biochemical analyses. For live cell imaging, siRNA-treated cells were transfected in fluorodish on day 5 with respective plasmids and imaged at day 6. For siRNA transfections (A549 cells), cells were grown in 6-well plates to ∼50 % confluency the day before transfection, and transfected (100 pmol/ well) using DharmaFECT (Dharmacon) for 48h, and infected or mock-infected (i.e., non-infected; similar protocols as for infection, but without virus addition) with PR8 at MOI 3 for 8h.

#### Infection with pathogens

For bacteria infection, 1.5 × 10^5^ HeLa cells were seeded in 12-well plates before transfection with siRNA by Lipofectamine RNAimax (Invitrogen) for 24 h following the manufacturer’s instructions. Transfected cells were infected with *C. trachomatis* serovar D stably transformed with the p2TK2-SW2 plasmid expressing mCherry (Agaisse and Derré, 2013) in triplicates at a multiplicity of infection (MOI) of 0.2, with a 30 min centrifugation step at 270x*g* at 37 °C. For viral infection, reverse-genetics derived A/Puerto Rico/8/34 (PR8 WT; H1N1) was used as a model virus and titrated according to reference (Vale-Costa et al., 2016). Virus infections were performed at a MOI of 3 for 8h. After 45 min, cells were overlaid with DMEM containing 10 % fetal bovine serum (Gibco, Life Technologies, 10500-064) and 1 % penicillin / streptomycin mix (Biowest, L0022-100).

### Plasmids

The KIF13A-YFP and mCherry-RAB11A were obtained by recombination by using the GATEWAY cloning system (Invitrogen) (Delevoye et al., 2014); GFP-RAB11A was a kind gift from Dr. Bruno Goud (Institut Curie, Paris, France); mCherry-KIF13B plasmid was a kind gift from Pr. Kaori Horiguchi Yamada (University of Illinois, Chicago, USA); plasmid coding for the PI4P sensor (pEGFP-SidC^609-776^) was previously described (Domingues et al., 2016); plasmid coding for the PI3P sensor (pEGFP-2x-FYVE domain) was kind gift from Pr. Harald Stenmark (Department of Molecular Cell Biology, Institute of Cancer Research, The Norwegian Radium Hospital, Oslo, Norway). The plasmids coding for the following protein products were obtained from Addgene: non-fluorescent FRB-ECFP(W66A)-Giantin [# 67903, pEGFP-C1; Pr. Dorus Gadella (van Unen et al., 2015)]; mRFP- PJ-Sac -FKBP [Pseudojanin (PJ)-Sac is a mutated chimeric phosphatase which dephosphorylates PI4P to produce PI] [# 38000, pmRFP-C1; Dr. Robin Irvine; (Hammond et al., 2012)]; iRFP-FRB- RAB5 [# 51612, pEGFP-C1; Pr. Tamas Balla (Hammond et al., 2014)]. The BLOC-1 expression plasmids (pST39-BLOC-1) is described in (Lee et al., 2012). The GST expression plasmid (pGST-parallel-1) is described in (Sheffield et al., 1999)

### siRNAs

The sense strand of the following siRNAs was synthesized by Qiagen: 1) Control: 5’- AATTCTCCGAACGTGTCACGT-3’; 2) PI4KIIα: smartpool (Flexitube siRNA, GS55361); 3) PI4KIIβ: 5’-CAGAGTACTGGCCTTGTTCAA-3’ (PI4KIIβ#7) and 5’- TCGGATTGTCCACCTGAGCAA-3’ (PI4KIIβ#9); 4) PI4KIIIβ: smartpool (Flexitube siRNA, GS5298); 5) BLOC-1 (individual siRNAs against the following BLOC-1 subunits): 5’- AATGCTGGATTCGGGAATTTA-3’ (snapin), 5’-ATGGTCCATGTTAAATGTAAA-3’ (muted), 5’-AAAAGTGCATGTACGGGAAAT-3’ (Pallidin#1), 5’-AACTGCAGCAGAAGAGGCAAA-3’ (Pallidin#2); 6) KIF13A: 5’-CTGGCGGGTAGCGAAAGAGTA-3’ (KIF13A#2), 5’-CCGCAACAACTTGGTAGGAAA-3’ (KIF13A#3). For all PI4KII knockdown experiments, expressions of PI4KIIα and of PI4KIIβ were extinguished by using a mix of individual siRNAs targeting PI4KIIα or PI4KIIβ.

### Antibodies and other reagents

#### Primary antibodies

sheep polyclonal anti-TGN46 (Biorad; 1:200, IFM); mouse monoclonal anti-β-actin [Sigma A5316; 1:1000, immunoblotting (IB)]; rabbit polyclonal anti-PI4KIIβ (1:1000, IB) and -PI4KIIα (1:1000, IB) were generous gifts from Pr. Pietro De Camilli (Yale School of Medicine, New Haven, USA); rabbit polyclonal anti-β-tubulin (Abcam ab6046; 1:1000, IB); rabbit polyclonal anti-Pallidin (Proteintech,10891-1-AP; 1:1000, IB); rabbit polyclonal anit-RAB11A (Proteintech, 15903-1-AP; 1:200, IFM); mouse monoclonal anti-NP (Abcam, 20343; 1:1000, IF); rabbit polyclonal anti-NP (1:1000), -PB1 (v19/6, 1:500), -PB2 (2N580, 1:500), -PA (1:500) and -NS1 (v29, 1:500) for IB were all kindly provided by Prof. Paul Digard (Roslin Institute, UK); mouse monoclonal anti-M2 (Abcam, 5416, clone 14C2; 1:500, IB); mouse monoclonal anti-actin (Sigma, A5441, 1:1000, IB); goat anti-M1 (Abcam, 20910; 1:500, IB); goat polyclonal anti-GST (GE healthcare, 27-4577-01; 1:2000, IB); rabbit polyclonal anti-Cap1 [1 μg/ml, prepared as described in (Hamaoui et al., 2020)].

#### Secondary antibodies

HRP-conjugated goat anti–mouse and anti–rabbit, and goat anti-donkey (Abcam, ab6721 and ab6789; 1:10000, and Thermo Fisher Scientific, A15999; 1:2000, IB), or donkey anti-goat (thermo Fisher Scientific, A15999; 1:2000, IB in PIP binding assay); fluorescent antibodies from IRDye range (LI-COR Biosciences; 1:10000, IB); Alexa Fluor (AF)-488, -555 or -647-conjugated anti-rabbit, or anti-sheep, or anti-mouse (Invitrogen; 1:200, IFM).

#### Other reagents

AF-488 (1:500) labelled-Tf and AF-647 labelled WGA (1:200) were from Invitrogen (Thermo Fisher Scientific); siR-Tubulin (1:1000) was from Spirochrome. PAO (PhenylArsineOxide), wortmannin, and rapamycin were from Sigma Aldrich and stock solutions were dissolved in dimethyl sulfoxide (DMSO; Sigma Aldrich). The following reagents for *in vitro* studies were from Avanti Polar Lipids: Egg phosphatidylcholine (EPC), brain phosphatidylserine (bPS), brain phosphatidylinositol-4-phosphate PI4P, 18:1 PI3P, C24:1 Galactosyl(β) Ceramide, DOPS (1,2-dioleoyl-*sn*-glycero-3-phospho-L-serine).

### Drug treatments

Cells were incubated 30 min with PAO (300 – 600 nM) or with Wortmannin (10 mM), or 20 min with 1 mM of Rapamycin (1 mM) prior to analyses by live cell imaging. Control cells were treated with the same volume of DMSO alone.

### Preparation of purified BLOC-1

#### Preparation of BLOC-1 for in vitro experiments using membrane systems

BLOC-1 was prepared according to (Lee et al., 2012) with some modifications. The pST39-BLOC-1 plasmid encoding recombinant BLOC-1 was transformed into BL21gold(DE3)plysS cells (230134, Agilent). Several colonies from the plate were inoculated into a starter culture of 10 ml LB supplemented with 34 μg/ml chloramphenicol and 100 μg/ml ampicillin that was grown overnight at 37 °C with moderate shaking. Next day the starter culture was transferred to 1 liter Terrific Broth medium supplemented with antibiotics and grown at 37 °C, 200 rpm for ∼7h. Cultures were induced with 0.8 mM IPTG and grown overnight at 16 °C, 200 rpm. Cells were centrifuged at 4 °C, 6000xg for 15 min and bacterial pellets were resuspended in buffer containing 50 mM Tris (pH 7.4), 300 mM NaCl, 5 mM β-mercaptoethanol supplemented with lysozyme, DNase I, and complete-EDTA free protease inhibitor tablets (1183617001, Roche) and incubated at 4 °C with rotation for 30 min. The sample was further sonicated, and the soluble fraction was separated by centrifugation at 4 °C, 35,267xg for 45 minutes. The supernatant was loaded on a gravity chromatography column preloaded with glutathione-Sepharose 4B (17-0756-05, GE Healthcare) and incubated for 2 h at 4 °C with gentle rotation in batch mode. Following incubation, the column was extensively washed with buffer (50 mM Tris (pH 7.4), 300 mM NaCl, and 5 mM β-mercaptoethanol) and on-column cleavage of the GST and 6xHis tags using tobacco etch virus (His_6_-TEV) S219V protease (∼0.5mg/ L) was done overnight at 4 °C. The flow-through fraction containing the cleaved BLOC-1 was collected and loaded onto a Ni-NTA column. Flow-through material was collected, concentrated, and further purified by size exclusion chromatography on a Superose 6 10/300 column in buffer containing 50 mM Tris (pH 8.0), 0.4 M NaCl, 2 mM DTT, 5 % glycerol and 1 mM EDTA. Fractions containing BLOC-1 were pooled together, aliquoted, flash frozen in liquid nitrogen, and stored at -80 °C. The integrity of the complex was analyzed by SDS-PAGE.

#### Preparation of GST and BLOC-1-GST

Plasmids encoding GST and BLOC-1-GST were expressed and purified as described above except that after the binding of the protein to glutathione sepharose and following the washing step, the bound GST proteins were eluted from the beads with elution buffer containing 50 mM Tris (pH 8.0) and 10 mM glutathione. 21 nM of GST or BLOC-1-GST were used for the PIP array assays. Assays were carried out as described in the following section.

### PIP array assay

PIP array (P-6100, Echelon Biosciences) binding assays were carried out according to the manufacturer’s protocol. Briefly, PIP membranes were blocked in PBS-T (PBS supplemented with 0.1 % v/v Tween-20) supplemented with 3 % fatty acid free BSA for 1 hour at room temperature with gentle shaking. Following blocking, membranes were incubated with the proteins that were diluted in blocking buffer and incubated for 1 hour with gentle agitation. Membranes were washed 3 times in 5 ml PBS-T for 5 min and then incubated with anti-GST antibody for 1 h at room temperature with gentle agitation.

Following incubation, membranes were washed 3 times in 5 ml PBS-T for 5 minutes and incubated with HRP antibodies. Following incubation, membranes were washed 3 times in 5 ml PBS-T for 5 min and an additional final wash of PBS. Membranes were blotted with Pierce™ ECL Western Blotting Substrate (cat# 185698) and signal was detected with Bio-Rad Chemidoc MP imaging system.

### In vitro experiments using membrane systems

#### Preparation of lipid nanotubes and binding assay with BLOC-1

Galcer/ EPC nanotubes doped or not with PIxP were prepared according to (Wilson-Kubalek et al., 1998). Briefly, a mixture of Galcer, EPC, and bPI4P or 18:1 PI3P was dried under vacuum. Galcer/ EPC (80/ 15 mol/mol), Galcer/ EPC/ bPI4P (80/ 15/ 5 mol/mol), Galcer/ EPC/ 18:1 PI3P (80/ 15/ 5 mol/mol) tubes were formed at room temperature after resuspension of the dried film at 5 mg/ml total lipid concentration in 50 mM Hepes pH 7.4 and 120 mM potassium acetate (HK buffer) followed by 5 cycles of 10 min vortex, 2 min at 40 °C. For experiments, tubes were aliquoted and stored at -20 °C. Tubes were diluted in HK buffer at 50 μM and BLOC-1 was added at a lipid/ BLOC-1 ratio of 300 mol/mol. After 5 min incubation, sample was used for EM analysis.

#### Preparation of lipid suspension and binding assay with BLOC-1

Dried films made of EPC/ PI4P (95/ 5 mol/mol) was made by evaporation of organic solution under vacuum for 1 h. The lipid film was resuspended in HK buffer at 1 mM while vortexing. This led to the formation of various population of vesicles and tubules. For cryo-EM, the lipid suspension was diluted to 50 μM and BLOC-1 was added at lipid/ BLOC-1 ratio of 300 mol/mol and incubated 5 min at RT before grid preparation for cryo-EM.

#### Giant Unilamellar Vesicle (GUV) preparation

GUVs were generated using a lipid mixture composed of EPC/ 0.5 mole% TR ceramide/ 10 mole% DOPS/ 5 mole% PI4P. BODIPY^TM^ TR ceramide (ThermoFisher) ceramide allows us to visualize GUVs by confocal fluorescence microscopy. The inner buffer used to generate GUVs is 50 mM NaCl, 20 mM sucrose and 20 mM Tris pH 7.5. GUVs were prepared by using the polyvinyl alcohol (PVA) gel-assisted vesicle formation method as previously described in (Weinberger et al., 2013). Briefly, a PVA gel solution (5 %, w/w, dissolved in 280 mM sucrose and 20 mM Tris, pH 7.5) warmed up to 50 °C was spread on clean coverslips (20mm × 20mm). Before use, the coverslips were washed once with ethanol and twice with ddH_2_O. For experiments, the PVA-coated coverslips were incubated at 50 °C for 30 min, and then approximately 5 μL of the lipid mixture was spread on the PVA-coated coverslips. The coverslips were then placed under vacuum for 30 min at room temperature, and then in a petri dish. Approximately 500 μL of the inner buffer was pipetted on top of the coverslips. The coverslips were kept at RT for at least 45 min, allowing GUVs to grow. Once done, we gently “ticked” the bottom of the petri dish to detach GUVs from the PVA gel. GUVs were collected using a 1 mL pipette tip with its tip cut to prevent breaking GUVs.

### Electron Microscopy

#### Negatively stained images of BLOC-1

BLOC-1 (25 μg/ml) was deposited on a carbon-coated EM grid previously glow-discharged and negatively stained with 2 % uranyl acetate in water. Negatively stained images of BLOC-1 were acquired as described below.

#### Cryo-EM experiments, imaging and analysis

Lacey carbon 300 mesh grids (Ted Pella, USA) were used in all cryo-EM experiments. In all experiments, blotting was carried out on the opposite side from the liquid drop and samples were plunge frozen in liquid ethane (EMGP, Leica, Germany). Cryo-EM images were acquired with a Tecnai G2 (Thermofisher, USA) Lab6 microscope operated at 200 kV and equipped with a 4k x 4k CMOS camera (F416, TVIPS). Image acquisition was performed under low dose conditions of 10 e^-^/Å^2^ at a magnification of 50,000 or 6500 with a pixel size of 0.21 or 1.6 nm, respectively. The diameter of the tubes depicted in Figure 2 G were measured from 65 tubes in 17 images from 2 experiments. Diameters were measured every 60 nm along a tube leading to a total of 408 values.

#### High Pressure Freezing, sample preparation, and conventional EM imaging

HeLa cells were seeded on carbonated sapphire discs and grown for 2 days prior to transfection with either control or PI4KII siRNAs. After two consecutive siRNA transfections, cells were immobilized by HPF using either a HPM 100 (Leica Microssystems, Germany) or a HPM Live µ (CryoCapCell, France), and freeze substituted in anhydrous acetone containing 1 % OsO_4_/ 2% H_2_O for 64h using the Automatic Freeze Substitution unit (AFS; Leica Microsystems). The samples were included in Epon. Ultrathin sections were contrasted with uranylacetate/ H2O and Reynold’s lead citrate solution. Electron micrographs were acquired using a Transmission Electron Microscope (Tecnai Spirit; Thermofisher, Eindhoven, The Netherlands) operated at 80kV, and equipped with a 4k CCD camera (Quemesa, EMSIS, Muenster, Germany).

### Fluorescence Microscopy

#### Live-cell

Hela cells were grown on fluorodish and imaged in their respective media supplemented with 20 mM HEPES (GIBCO, ThermoFisher Sicentific).

#### Sample preparation of GUVs and fluorescent imaging

Experimental chambers were assembled by sandwiching two coverslips using parafilm®. Before use, the chambers were passivated with a β-casein solution at a concentration of 5 mg/ml for at least 5 min at room temperature. GUVs were incubated with BLOC-1 at a final concentration of 0.2 μM in the experimental chambers for at least 15 min at RT before observation. To mix GUVs with BLOC-1, we first added 8 μL of the GUVs in 15 μL of an “outer buffer” (60 mM NaCl, 20 mM Tris, pH 7.5), following by adding 2 μl of BLOC-1 (2.5 μM in stock). Samples were observed using a spinning disk confocal microscope, Nikon eclipse Ti-E equipped with Yokogawa CSU-X1 confocal head, 100X CFI Plan Apo VC objective (Nikon) and a CMOS camera, Prime 95B (Photometrics). After formation of tubules, sample was pipetted off and flash frozen for cryo-EM expriments.

#### Immunofluorescence (IF)

Hela cells grown on coverslips were first fixed with 4 % PFA in PBS for 15 min at RT and then subsequently incbubated for 5 min each with PBS/ 2 % glycine and PBS/ 0.2 % BSA (blocking buffer) and PBS/ 0.2 % BSA/ 0.1 % Saponin (permeabilization buffer). The cells were further washed three times with permeabilization buffer and incubated with AF-conjugated secondary antibodies diluted in permeabilization buffer before mounting with Prolong Gold Antifade (Invitrogen). For viral infection studies, A549 cells were similarly fixed with 4% paraformaldehyde, permeabilized with 0.2% TritonX and stained in PBS/1% PBS and processed similary as described above. For bacterial inclusion studies, twenty hours after infection, HeLa cells were fixed and permeabilized for immunofluorescence using 0.05 % saponin, 5 mg/ ml BSA in PBS.

#### Fluorescence Imaging

HeLa cells were imaged on an inverted Eclipse Ti-E (Nikon) + Spinning disk CSU-X1 (Yokogawa) integrated in Metamorph software (Molecular devices, version 7.8.13) with an EMCCD camera (iXon 897; Andor Technology), a 100× 1.4 NA Plan- Apo objective lens and Z-images were taken at 0.2 mm with the piezoelectric motor (Nano z100, Mad City Lab). For imaging of virus-infected cells, single optical sections were imaged with a SP5 live confocal microscope (Leica). Images were post-processed using ImageJ/ FIJI (NIH).

#### Super-resolution (SR) microscopy

images were acquired on a spinning-disk system equipped with a super-resolution module (Live-SR; Gataca Systems, France) leading to super-resolution spinning-disk confocal microscopy using optical photon reassignment [as described before in (Azuma and Kei, 2015)]. This method allows combining the doubling resolution together with the physical optical sectioning of confocal microscopy. The maximum resolution is 128 nm with a pixel size in SR mode of 64 nm.

### Tf recycling assay

Cells starved for 30 min in serum-free medium supplemented with HEPES (20 mM, pH 7.4; Invitrogen) and 0.5 % (w/v) BSA medium were pulsed for 10 min with Tf-AF-488. Cells were then quickly incubated with cold acid buffer (0,5 M Glycine, pH 2.2) to dissociate surface-bound Tf, extensively washed with PBS, and chased at 37 °C with complete media (containing 20 mM HEPES) for various time points prior to chemical fixation with 4 % paraformaldehyde (Thermo Fisher Scientific) and processing by FM.

### Immunoblotting

Hela cells were trypsinized, pelleted, washed with cold PBS and incubated in cold lysis buffer [50 mM Tris, 150 mM NaCl, 0.1 % Triton X-100, 10mM EDTA, pH 7.2, supplemented with protease inhibitor cocktail (Roche)]. After the cell lysis, post-nuclear supernatants were obtained by centrifugation. Cell lysates (∼5 μg) were incubated with sample buffer, boiled for 5-10 min before the loading on 4-12 % Bis-Tris gels (Nu-PAGE, Invitrogen) and then transferred onto nitrocellulose membranes (GE Healthcare). The membranes were blocked in PBS (with 0.5 % Tween 20, 5 % non-fat dried milk) for 30 min and incubated with respective primary for 8-12 hours and 30 min with secondary antibodies diluted in PBS/ 0.1% Tween 20. Membranes were washed with PBS/ 0.1% Tween 20 twice for 30 min and developed using ECL SuperSignal West (Pico or Dura, Thermo Fisher Scientific) and visualized by using X-ray films (Kodak developer). For virus-infected cell lysates, IB was performed as above and imaged using a LI-COR Biosciences Odyssey near-infrared platform.

### Quantitative real-time reverse-transcription PCR (RT-qPCR)

RNA was extracted from samples in NZYol (NZYtech, MB18501) using Direct-zol RNA minipreps (Zymo Research, R2052). Reverse transcription (RT) was performed using the NZY MuLV first strand cDNA synthesis kit (NZYTech, MB17302). Real-time RT-PCR to detect GAPDH, PI4KIIα, PI4KIIβ and PI4KIIIβ was prepared in 384-well, white, thin walled PCR Plates (ABgene 12164142) using SYBR Green Supermix (Biorad, 172-5124), 10% (v/v) of cDNA and 0.4 µM of each primer. The reaction was performed on an ABI QuantStudio-384 (Applied Biosciences) under the following PCR conditions: Cycle 1 (1 repeat): 95 °C for 2 min; Cycle 2 (40 repeats): 95 °C for 5 s and 60 °C for 30 s; Cycle 3: 95 °C for 5 s and melt curve 65 °C to 95 °C (increment 0.05 °C each 5 s). Data were analyzed using QuantStudioTM Real Time PCR software (v1.1., Applied Biosciences). Primer sequences used for real-time RT-qPCR were the following: 1) GAPDH — Fw: 5’- CTCTGCTCCTCCTGTTCGAC-3’, Rv: 5’-ACCAAATCCGTTGACTCCGAC-3’; 2) PI4KIIα — Fw: 5’-TCTTTCCCGAGCGCATCTAC-3’, Rv: 5’-GCAGCCACTTGGTCCACTTA-3’; 3) PI4KIIβ — Fw: 5’-AAGCGGGTGCCTATCTTGTG-3’, Rv: 5’-TTTTGCACGGTCAATCGCAT- 3’; 4) PI4KIIIβ — Fw: 5’-TCAGCAGCAACCTGAAACGA-3’, Rv: 5’- CAGTCGAACAGGGGAACTGA-3’.

### Image analyses and quantifications

#### KIF13 Tubule quantification

Quantification of the average number of cells with at least one KIF13A^+^ tubule was done manually (**Fig. 3**, **D** and **H**; and **Fig. S3 D**). Quantification of the average length of KIF13A^+^ tubules was carried out using ImageJ/Fiji as described previously (Delevoye et al., 2016; Dennis et al., 2015). In short, live cell fluorescent image was converted into binary and skeletonized. The resulting skeletonized image was then analyzed using the integrated plugin ‘AnalyzeSkeleton’. The skeletonized image was manually compared with the raw image to exclude false-positive tubules from the analysis. The length and number of tubules were determined after applying a threshold of 8 pixels (1.28 μm). Only tubules longer than 8 px were included in the analysis (**Fig. 3 E**; and **Fig. 4**, **D-E**).

#### Maximum length, width, and neck width of endosomal nascent tubules

EM images were analysed using the iTEM software that was used to manually draw lines along the long axes of length, width or neck of the endosomal nascent tubules. The obtained values were collected in nm and statistical analyses were performed using Graphpad Prism software (version 9) (**Fig. 5 D**; and **Fig. S4**, **B** and **D**).

#### Measurement of fluorescence intensity

Fluorescence intensity of Tf labeling was measured using the “Measure” function of imageJ/Fiji on average z-projections from the raw images. For each condition, the intensities were normalized to their respective mean at 0 min of chase (**Fig. S5**, **B-C**).

#### Line scan analysis

Hela cells co-expressing mCherry-KIF13B and SidC-EGFP or EGFP-FYVE were analysed by live SR-FM. Using the image of the mCherry channel, a line of 30 pixels was drawn crossing the mCh-KIF13B tubule and the fluorescence intensities of the mCherry and of the GFP was measured per pixel using the “Plot Profile” function of imagej/Fiji and normalized to the maximum value (**Fig. 1 D**).

#### Image processing

Images were treated with Clarify.AI module (deep neural network trained to recognize and remove out-of-focus signal on each plane) proposed by Nikon NIS-Elements software in the NIS.AI (**Fig. 1 A**, top panel; and **Fig. 6 D**).

#### Immunoblot quantification

Protein expression levels were quantified using imagej/Fiji software and intensities were normalized to the respective bands of the loading control. The quantification was done on 3 independent experiments.

#### Bacterial inclusion analyses

5 – 10 pictures were taken randomly for each coverslips using the imagej/Fiji software, the scale was set from pixels to micrometers and the length in µm of budding tubules was measured by using ImageJ/Fiji tracing tool and number of tubules were counted using the multi-point tool. We calculated the mean number of tubules and the average tubular length per cell for each condition. Each condition was analyzed in a blind fashion from three individual coverslips with a minimum of 50 inclusions per coverslips.

#### Statistical analysis

Statistical data are represented as mean ± SEM. Graphpad (Prism 7 or 9) was used for all statistical analyses. Either unpaired two-tailed Student’s t-test or ordinary Anova test was used, as indicated in the Figure legends. The significant differences noted are between the control and test condition, and only p< 0.5 was considered significant ( *p < 0.05; **p < 0.01; ***p< 0.001 ****p < 0.0001).

## Figure Legends

**Figure S1. PI4P and KIF13 do not associate with early sorting endosomes**

**A** and **B**) Live imaging frame of a Hela cell co-expressing (**A**) mCh-KIF13B (red) and iRFP- RAB5 (pseudocolored in green) or (**B**) GFP-coupled PIxP sensors (green) for either PI4P (SidC-GFP) or PI3P (GFP-2x-FYVE) with iRFP-RAB5 (pseudocolored in red). Magnified insets (4x) show that RAB5^+^ structures overlap with PI3P (**B**, bottom panels; arrowheads) but do not overlap with KIF13B^+^ RE tubules (**A**; arrowheads) or PI4P (**B**, top panels; arroweads). **C**) IFM of fixed Hela cells expressing the PI4P sensor (GFP-SidC, green) and labeled for markers (red) of the TGN (TGN46; top panel) or of the plasma membrane (fluorescent-conjugated Wheat Germ Agglutinin, WGA; bottom panel). Arrowheads, point areas of overlap. Cell periphery is delimited by yellow lines. Bars: (main panels) 10 µm; (magnified insets) 2 µm.

**Supplementary Figure 2. BLOC-1 binding to vesicles and formation of tubules**

**A**) Schematic of the polycistronic expression cassettes used for the expression and purification of recombinant BLOC-1 in pST39 (left panel) and SDS PAGE gel of the purified BLOC-1 (right panel). **B**) Lipid strip assay showing interaction of BLOC-1 with various PIxPs. **C**) Cryo-EM images of control GalCer/EPC nanotubes in absence of BLOC-1. **D**) Cryo-EM images of control GalCer/EPC nanotubes in presence of BLOC-1. No protein was bound to nanotubes (black arrowheads). **E**) Cryo-EM image of BLOC-1 bound to GalCer/ EPC/ 18:1 PI3P (80/ 15/ 5) nanotube (white arrowheads). **F**) Cryo-EM image of BLOC-1 bound to GalCer/ EPC / bPI4P (80/ 15/ 5) (white arrowheads). **G**) Maginified region of (F) showing dark dotted densities corresponding to BLOC-1 bound to PI4P^+^ nanotube. **H**) Cryo-EM image of a suspension of EPC/ bPS/ bPI4P (85/ 10/ 5 mol/mol) vesicles and tubes incubated with BLOC-1 (0.15 μM proteins and 50 μM lipids). BLOC-1 binds to tubules (white arrow), but not to large vesicles (black arrows). **I**) Cryo-EM image of tubules generated upon BLOC- 1 (white arrowhead) addition to GUVs of EPC/ bPS/ bPI4P (85/ 10/ 5 mol/mol) (see also Figure 2 **H**). Figures are representative of at least three independent experiments. Bars: (**C**- **I**) 25 nm.

**Figure S3. PI4KII enzymatic activity is required for the formation and stabilization of recycling endosomal tubules**

**A**) Live imaging frames of HeLa cells co-expressing KIF13A-YFP (top) and mCh-RAB11A (bottom) and treated with control (siCTRL) or PIK4II (siPI4KII) siRNAs. Note the tubular KIF13^+^ tubular structures in siCTRL-treated cell and the vesicular structures in siPI4KII- treated cells. Insets are magnifications of boxed areas showing the co-distribution of KIF13A with RAB11A (arrows). **B**) Live imaging frame of siCTRL- or siPI4KII- treated HeLa cells expressing KIF13A-YFP (top) and incubated with the siR-Tubulin probe (bottom) to detect microtubules. **C**) Live imaging frame of HeLa cells expressing KIF13A-YFP and treated with DMSO vehicle (left), the PI3K inhibitor wortmannin (10 µM, middle), or the PI4K inhibitor PAO (300 nM, right). **D**) Quantification of the average percentage of treated cells as in C in which at least one KIF13A^+^-YFP tubule (n > 60 cells) was evident. **E**) Time lapse images of HeLa cells expressing KIF13A-YFP before (0 min) and 30 min after (30 min) addition of PAO (600 nM). Data represent the average of at least three independent experiments and presented as mean ± SEM *p < 0.05. ns, not significant. Bars: 10 μm.

**Figure S4. Depletion of PI4KIIs did not alter the neck and width of endosomal nascent tubules**

**A**) Schematic representation of the measurements (taken in Fig. 5D and Fig. S4, B-D) of the endosomal nascent tubules observed by EM. **B-D**) Quantification of the average maximum width (B), neck width (C) and length:width ratio (D) of the endosomal nascent tubules in siCTRL- (nascent tubules: width, n = 42; neck, n = 30, length:width, n = 40) and siPI4KII- (nascent tubules: width, n = 50; neck, n = 49, length:width, n = 46) treated HeLa cells. Data are the average of two independent experiments presented as the mean ± SEM (siCTRL, n = 8 cells; siPI4KII, n = 14 cells). ns, not-significant. B and C: two-tailed unpaired *t*-test.

**Figure S5. PI4KII expression is required for endosomal cargo recycling**

**A**) Fluorescence microscopy of siCTRL- (left panels) or siPI4KII- (right panels) treated Hela cells that were pulsed with Tf-A488 (green) and chased at the indicated time (0, 20, 40 min) before fixation and labeling of nuclei with DAPI (blue). Arrowheads point to the perinuclear accumulation of Tf^+^ endosomes in siPI4KII-treated cells during the chase. **B**) Quantification of the average fluorescence intensity of Tf per cell at time 0 min of chase (siCTRL, n = 50 cells; siPI4KII = 46 cells). **C**) Quantification of the average intensity of Tf per cell at different time points of chase (siCTRL cells: 0 min, n = 50; 20 min, n = 58; 40 min, n = 49; siPI4KII cells: 0 min, n = 46; 20 min, n = 56; 40 min, n = 46) normalized relative to the 0 time point. Cell periphery is delimited by yellow lines. Data are the average of three independent experiments presented as the mean ± SEM. ns, non-significant. ***p < 0.001; ****p < 0.0001. B and C, two-tailed unpaired *t*-test. Bars: 10 μm.

**Figure S6. PI4KII, BLOC-1, or KIF13A expressions are required for viral or bacterial infection**

A549 cells treated with control (CTRL) or PI4KII siRNAs for 48h before being mock-infected or infected 8h with PR8 virus, at a multiplicity of infection of 3. **A** and **B**) mRNA expression levels of PI4KIIα or PI4KIIβ determined in relation to GAPDH levels by real-time RT-qPCR. **C** and **D**) Immunoblot analyses of cells treated as in A show expression levels of viral proteins PB1 (86.5 kDa, C), PB2 (85.7 kDa, C) and PA (84.2 kDa, C), and NP (56.1 kDa, C), or M1 (27.8 kda, D) and NS1 (26.8 kDa, D), as well as actin (42 kDa, D) as a loading control. Actin, NS1 and M2 are colored in green channel, whereas M1 (11kDa, D) is colored in red. **E**) Viral protein levels determined by immunoblotting using antibodies against the indicated proteins. **F**) Immunoblotting of HeLa cell lysates treated with control (CTRL) or siRNA agains BLOC-1 subunits or KIF13A and probed for KIF13A (top), Pallidin (middle; BLOC-1 subunit) and β-Tubulin (loading control, bottom). **G** and **H**) HeLa cells treated with siCTRL or siRNA to KIF13A (KIF13) or the subunits of BLOC-1 (BLOC-1) were infected with Ctr D, and then analyzed 12 h later by IFM for Cap1. The average number (G) and the mean length (H) of Cap1^+^ tubules emanating from the bacterial inclusion. Statistical analysis (A and B) was performed using one-way ANOVA followed by Dunn’s multiple comparisons test (*p < 0.05; ***p < 0.001; n = 7 independent experiments).Data (G and H) are the average of three independent experiments presented as the mean ± SEM. E and F, two-tailed unpaired *t*-test. *p < 0.05; **p < 0.01; ***p < 0.001.

## Notes

### Competing Interest Statement

The authors have declared no competing interest.

