## Supplementary figures and images for "PI4P and BLOC-1 remodel endosomal membranes into tubules"

### Supplemental Figures

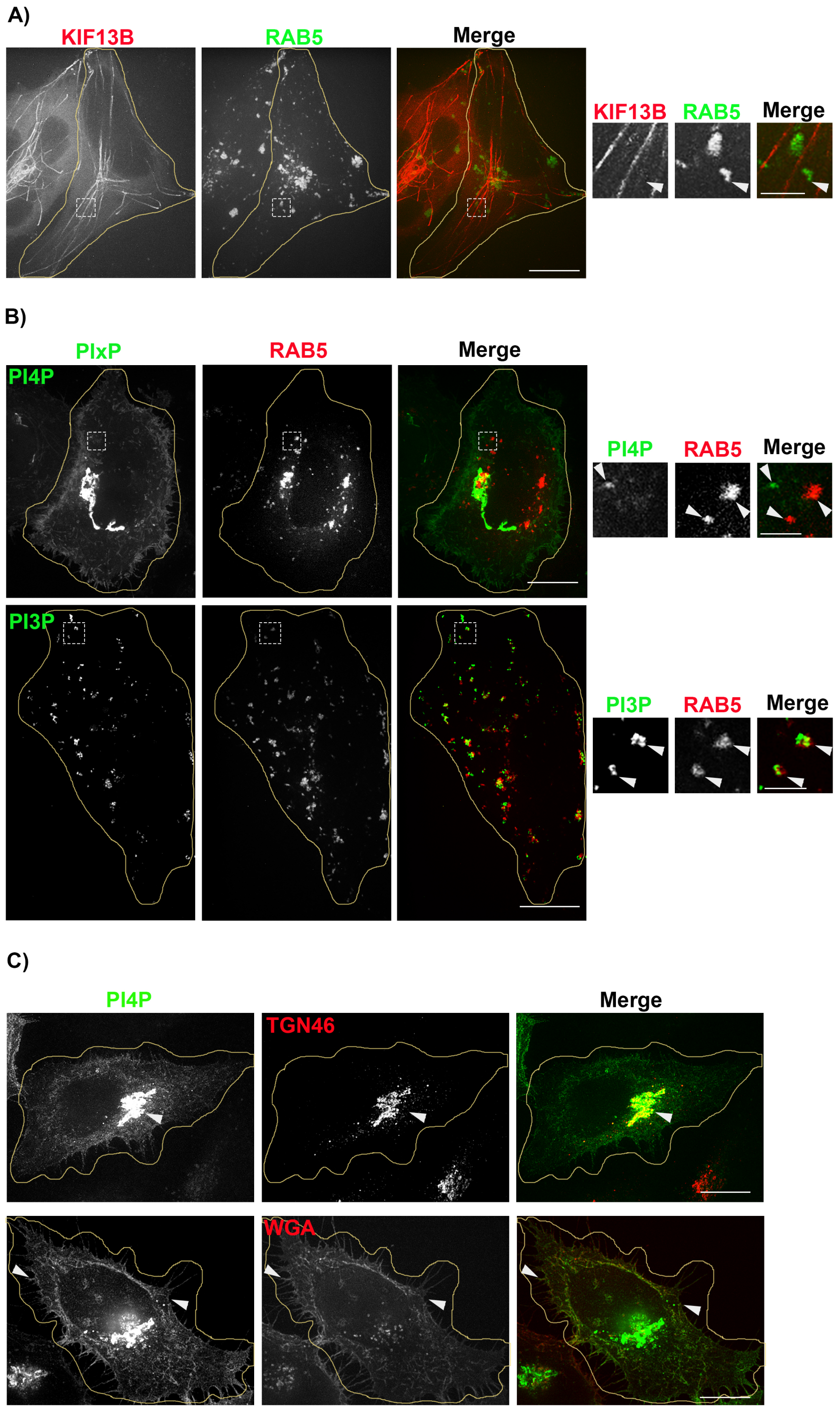

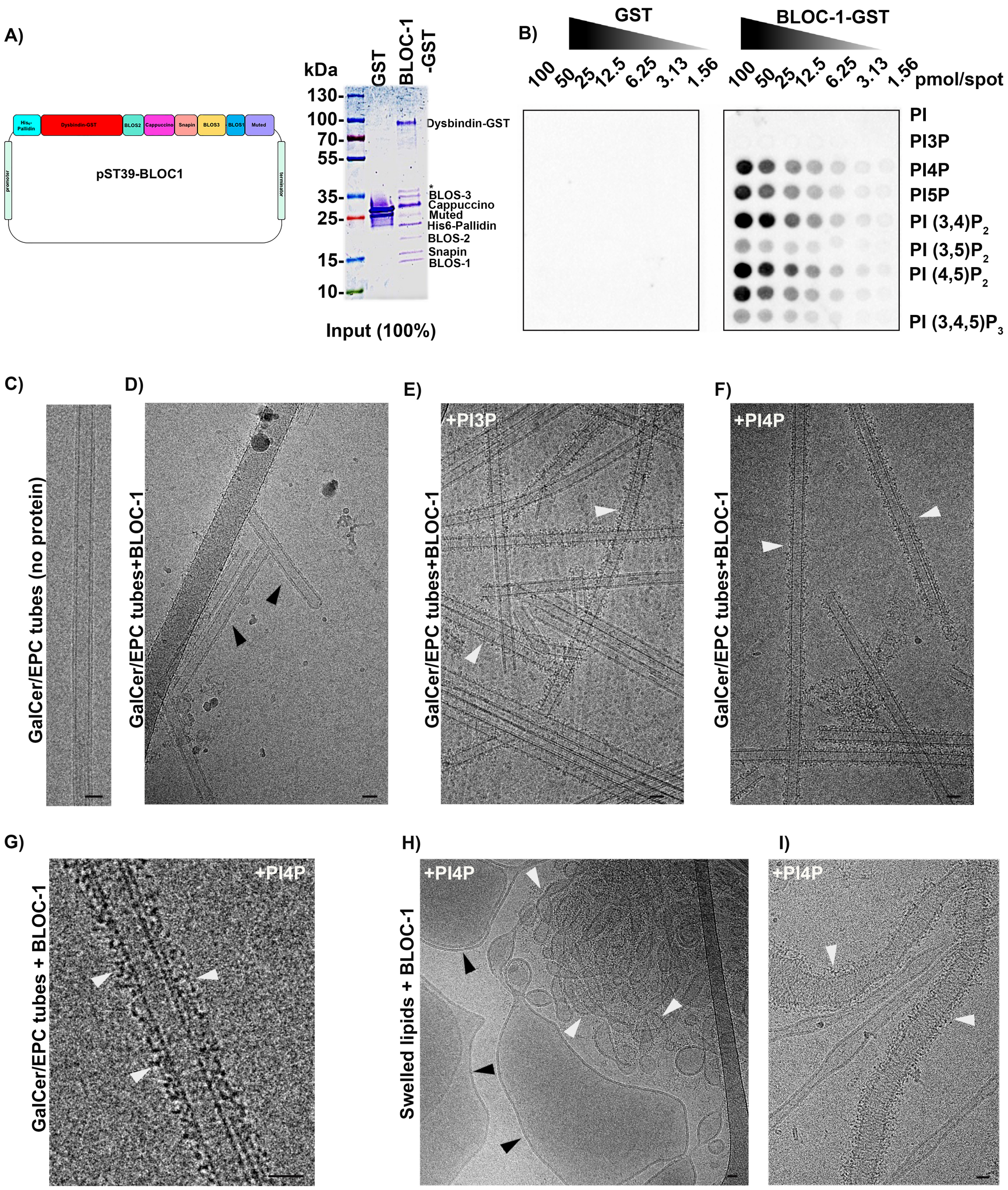

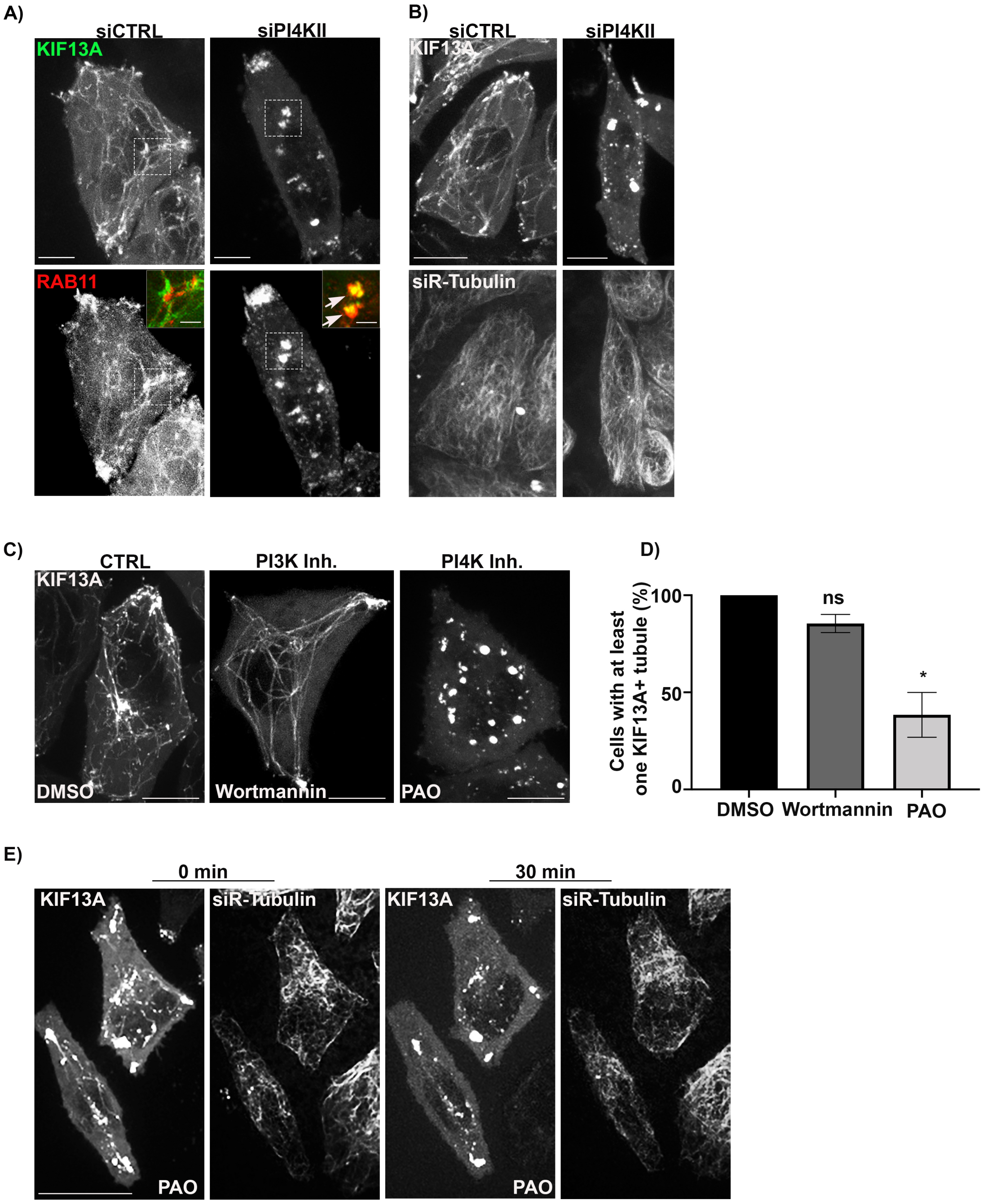

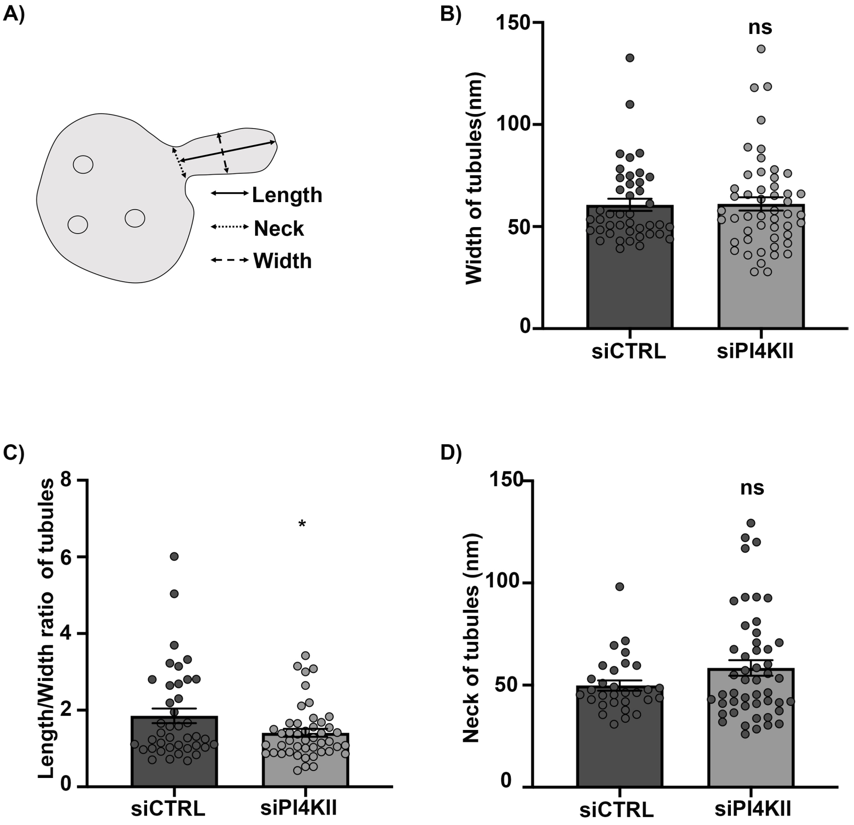

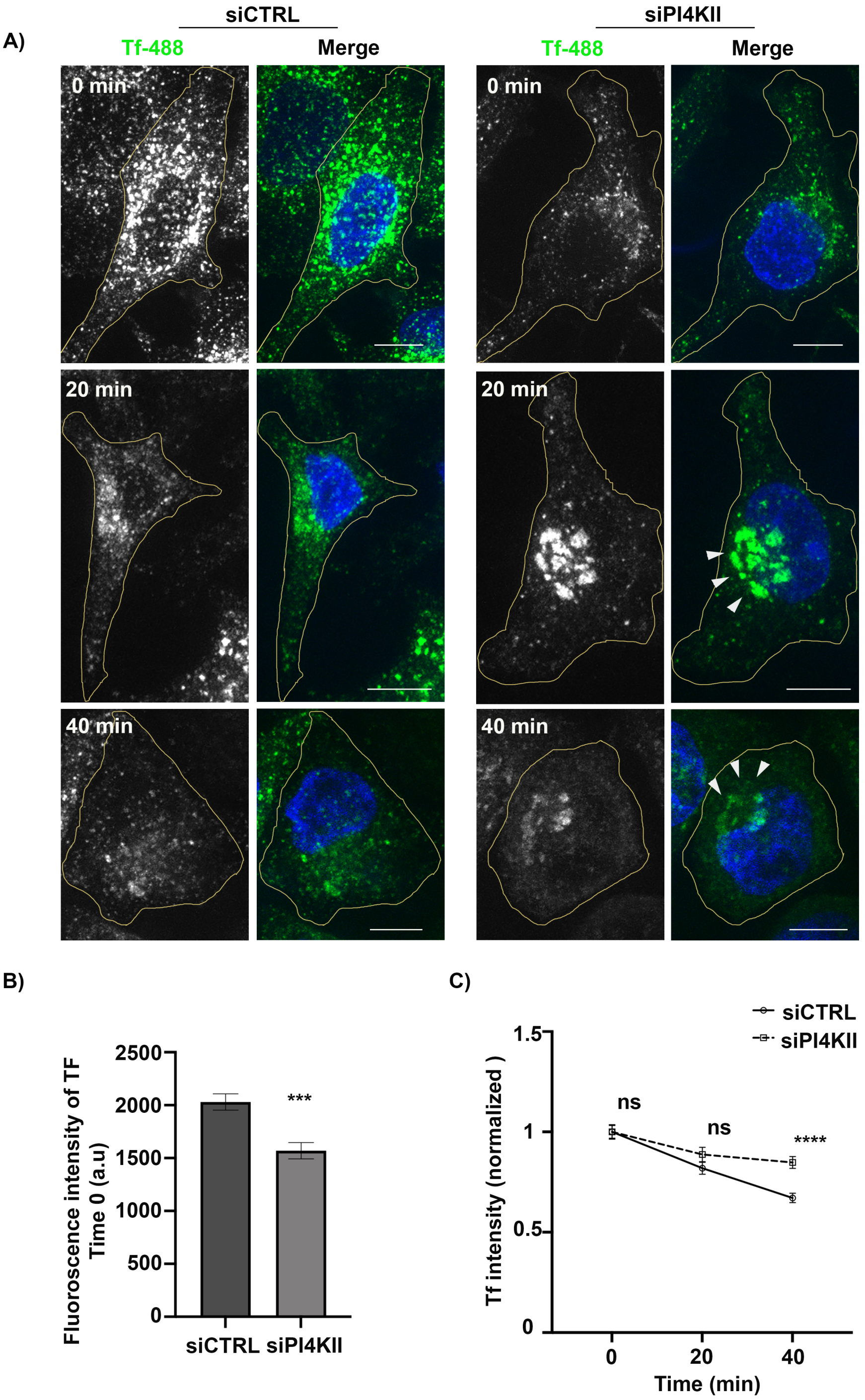

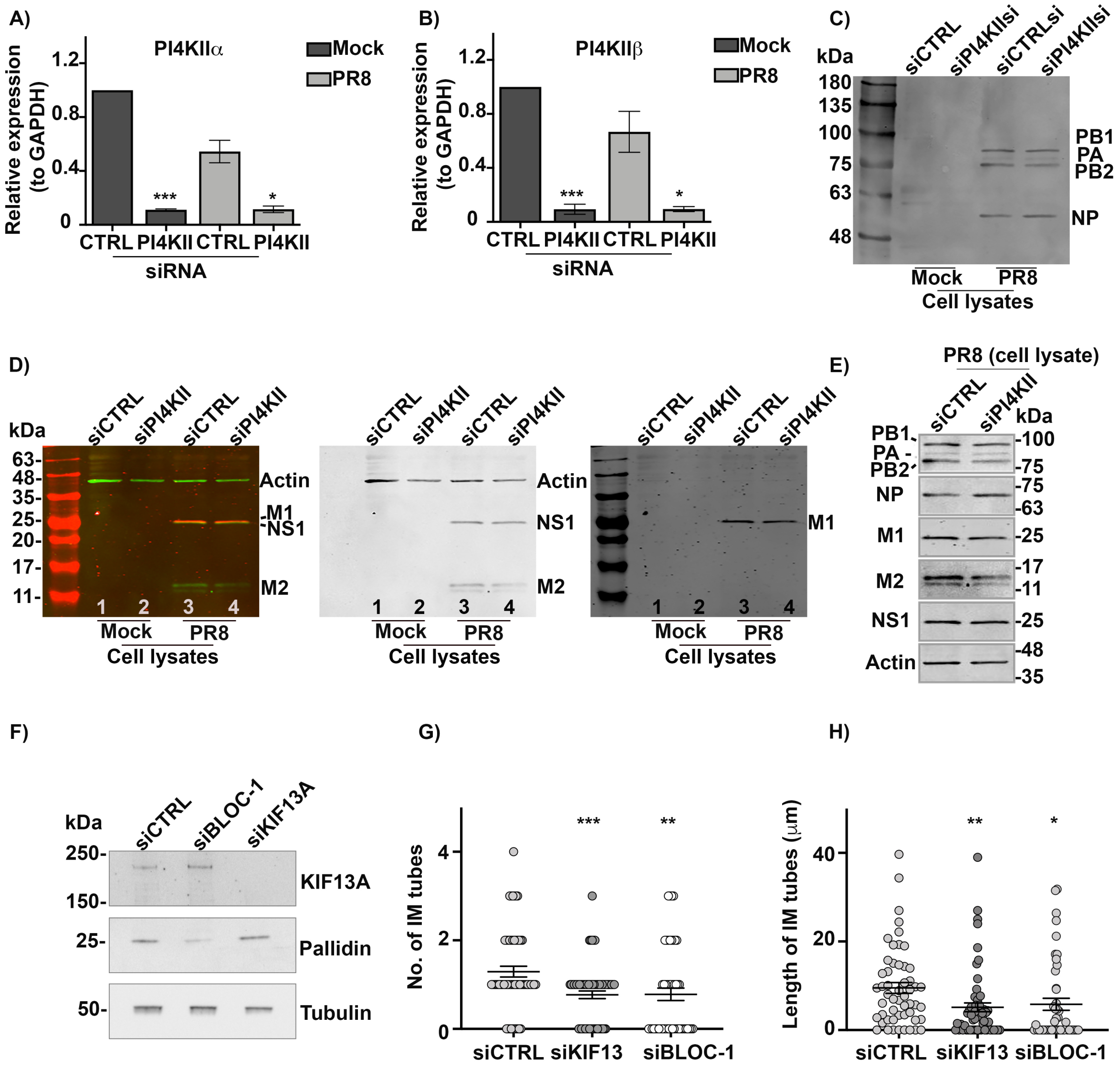
